# Identification of nuclear pore proteins at plasmodesmata: potential role in intercellular transport?

**DOI:** 10.1101/2024.09.02.610746

**Authors:** T. Moritz Schladt, Manuel Miras, Jona Obinna Ejike, Mathieu Pottier, Lin Xi, Andrea Restrepo-Escobar, Masayoshi Nakamura, Niklas Pütz, Sebastian Hänsch, Chen Gao, Julia Engelhorn, Marcel Dickmanns, Gwendolyn V. Davis, Ahan Dalal, Sven Gombos, Ronja Lange, Rüdiger Simon, Waltraud X. Schulze, Wolf B. Frommer

## Abstract

Plasmodesmata (PD) mediate intercellular exchange of small molecules, RNAs and proteins between plant cells with an apparent exclusion limit for passive non-specific transport, and transport of specific cargo mediated by mediators. PD and nuclear pore complexes (NPC) are nanometer sized micropores with strikingly similar properties. Cargo translocation through NPC is mediated by phase separating FG-nucleoporins (FG-NUP). Here, bioinformatics, proteomics and fluorescence imaging identified FG-NUPs at PD. Transient expression of GFP fusions at low and intermediate expression levels supported dual localization of 12 NUPs to NPC and PD. Structured illumination microscopy detected the transmembrane anchor NUP CPR5 close to orifices of PD. *cpr5* mutants showed reduced intercellular short-root (SHR) transport. However, transport defects cannot be excluded due to indirect effects in the mutants. Identification of FG-NUPs at PD is consistent with the recruitment of NUPs to form a PD pore gating complex consistent with phase separation domains as diffusion barriers at PD. Further analyses will be required to determine whether NUPs are *bona fide* PD components, or accumulate at PD in certain conditions, or may serve intermediate NPC storage.

## Introduction

Nutrient exchange and communication between cells are essential for multicellular organisms. Besides membrane transporters and cell surface receptors, metazoa, fungi and plants independently evolved cell-cell bridges. The green lineage developed unique structures, called plasmodesmata (PD), composed of modified cell wall domains that form plasma membrane-lined channels and that contain an ER-derived desmotubule providing continuity between the cytoplasm of neighboring cells (symplasm). Since PD are embedded into the cell wall and contain diverse membrane and soluble proteins, PD pose a major challenge for biochemical and cell biological analyses. Several studies obtained PD proteomes by enriching membrane proteins in cell wall fractions (Bayer et al., 2006; Brault et al., 2019; Fernandez-Calvino et al., 2011; Gombos et al., 2023). Genetic approaches failed so far to identify proteins that could fully explain the transport mechanism, possibly due to redundancy or lethality of mutants. Intercellular transport studies are also challenging since intact at minimum, a functional two cell system is required. Current models for PD translocation implicate a simple filter mechanism generated by spoke-like structures connecting the peripheral plasma membrane with the desmotubule, however the complex transport properties as well as inconsistencies such as the finding that sleeveless PD lacking an obvious path may put these models in question.

Key features of PD transport include the ability to translocate highly diverse cargo (from ions to macromolecules), an apparent size exclusion limit (which, depending of the type of PD, can reach 70 kDa, in funnel PD up to 120 kDa), and the ability to translocate specific cargo (Paultre et al., 2016; Ross-Elliott et al., 2017). Translocation through PD and the nuclear pore complex (NPC) share common features, namely both NPC and PD transport a similar spectrum of cargo species (ions, metabolites, proteins, and ribonucleoprotein complexes)(Ejike et al., 2025; Lee et al., 2000). For both NPC and PD, the ability for pore dilation has been described (Kragler et al., 1998; Zimmerli et al., 2021). Both NPC and PD constitute selective barriers, allowing ‘*passive passage of molecules’* with an apparent size exclusion limit (term used in animal literature; named ‘*non-targeted movement*’ for PD) and ‘*facilitated translocation of specific macromolecules’* (‘*targeted movement*’ for PD)(Crawford and Zambryski, 2001; Frey et al., 2018). Notably, the surface properties of cargo proteins play an important role for translocation, allowing even tetrameric GFP variants with compatible surface properties to pass (Frey et al., 2018). The core of the permeability barrier of the NPC appears to be a separated phase / condensate formed by disordered phenylalanine–glycine (FG) repeat-containing proteins, the FG-nucleoporins (Frey et al., 2018; Hoogenboom et al., 2021; Ng et al., 2021; Yu et al., 2023). These smart sieves mediate passage of smaller molecules without interaction with the FG repeats, while larger molecules or specific cargo can pass with the help of nuclear transport receptors that specifically interact with the FG repeats (Kehlenbach et al., 2023). As a shared structure among eukaryotes, the ultrastructure, function and transport mechanism of the NPC is well understood (Ejike et al., 2025; Mosalaganti et al., 2022). The human NPC, with about 1000 polypeptides, has an approx. diameter of 120 nm, a pore size of ∼43 nm and a height of 80 nm generated by membrane-anchors, three stacked rings (a sandwich of two outer and one inner ring), cytoplasmic filaments, and a nucleocytoplasmic basket (Lin and Hoelz, 2019). The human genome encodes twelve FG-nucleoporins (NUPs): NUP42, NUP50, NUP54, NUP58 (aka NUP45), NUP62, NUP98, NUP153, NUP214, POM121, POM121B, POM121C, and NUP358(Terry and Wente, 2009), while plant genomes encode ten FG-NUPs: NUP42 (aka CG1), NUP50a & NUP50b, NUP54, NUP58, NUP62, NUP98a & NUP98b, NUP136 and NUP214 (Chopra et al., 2019; Makarov et al., 2021). The majority of the FG-NUPs were conserved, with two FG-NUPs apparently lost in the green lineage (POM121, NUP358).

### FG-NUPs and other NUPs in PD fractions of *Physcomitrium patens* and localization to PD

Based on the striking similarities in transport properties, we hypothesized that PD transport may use permeability barriers similar as the NPC, that is, phase-separation generated by FG-repeat proteins. We thus explored whether the green lineage evolved a distinct set of FG-repeat containing proteins. A bioinformatic search for FG repeat-containing proteins in the genome of the moss *P. patens* revealed that 12 out of the 15 top-ranked FG-containing proteins were NUPs; the other three proteins were not considered candidates for a transport function (table S1; file S1; file S2 for Arabidopsis FG-containing proteins). An alternative hypothesis was that the PD permeability barrier is generated by NUPs derived by tandem gene duplications followed by differentiation into NPC and PD functions. However, phylogenetic analyses of Arabidopsis *NUPs* showed that the majority of NUPs are encoded by single genes (see also table S2 for overview of Arabidopsis *NUPs*)(Makarov et al., 2021; Zhang et al., 2020). Interestingly, however, the PD proteome from *P. patens* contained FG-NUP50a and two other NUPs, SEC13 and NUA (Gombos et al., 2023). Since PD cannot be purified to homogeneity, we cannot exclude that these NUPs represent contaminations. Previous work had indicated that *bona-fide* moss PD proteins can be targeted to PD when expressed in tobacco leaves (Gombos et al., 2023). To test whether FG-NUPs localize exclusively to the NPC or can also be targeted to PD, the subcellular localization of fluorescent protein (FP) fusions of three *P. patens* FG-NUPs (NUP58, NUP62 and NUP98) was analyzed by confocal microscopy after β-estradiol-controlled transient expression in *Nicotiana benthamiana*. A substantial number of fluorescent puncta derived from C-terminal mVenus fusions of the three FG-NUPs overlaid with fluorescent puncta of aniline blue marking callose at PD (Figure 1; Figure1-figure supplement 1). PD localization did not appear to be due to an FG-NUPs-specific mis-localization, since the inner ring linker NUP35, which connects the transmembrane interaction hub with the ALADIN-NDC1 core to NUP155, also localized to PD (Figure 1). Of note, the fluorescent puncta derived from mVenus fusions in the periphery did not exclusively localize to aniline blue labeled PD pit fields. Fluorescent mVenus and aniline blue derived puncta were also separately detected in locations without overlay. It is known that aniline blue does not detect all pit fields (Huang et al., 2022). Notably, transient expression in *N. benthamiana*, even though controlled by use of an estrogen-inducible system, can lead to overexpression; thus, the observed localization could either represent a physiologically relevant site for accumulation when under conditions of excess, or be due to accumulation in the vicinity of PD orifices.

**Figure 1.**
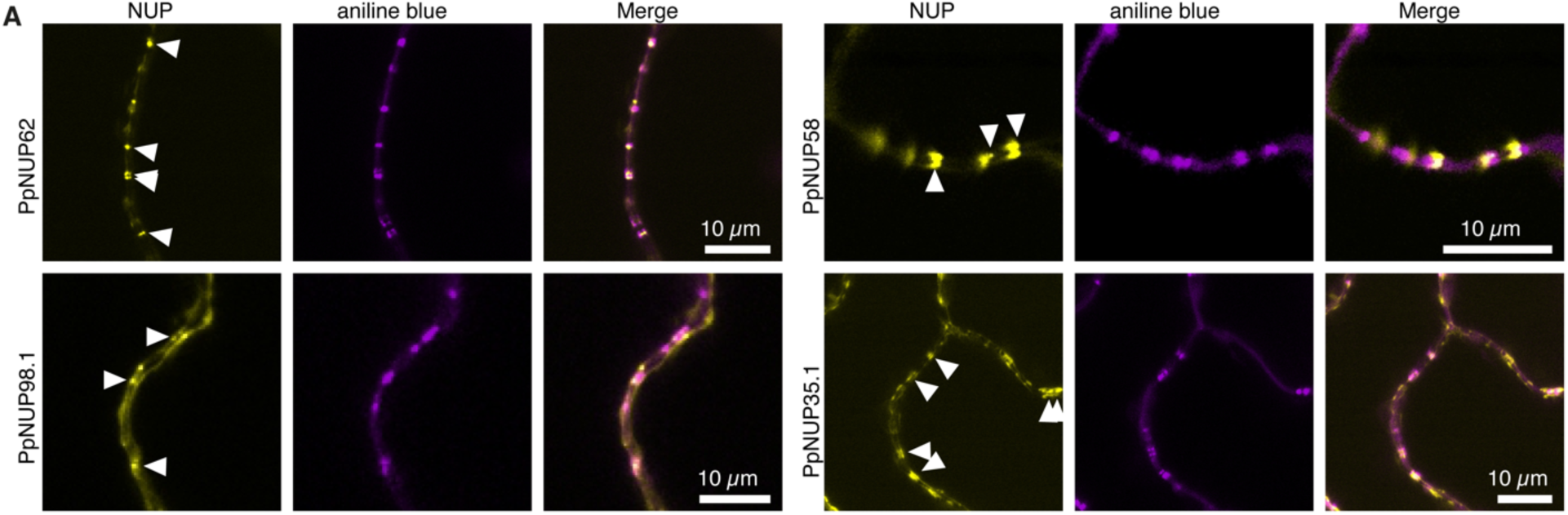
Identification of NUPs at PD. (**A)** Localization of NUPs from *Physcomitrium patens* at PD transiently expressed in *N. benthamiana* epidermal cells visualized by a single confocal optical section.

### FG-NUPs and other NUPs in PD fractions of Arabidopsis

To explore whether FG-NUPs may also be present in PD fractions from a vascular plant, we utilized established PD enrichment and data analysis pipelines to generate a PD proteome from Arabidopsis shoots (Gombos et al., 2023). This shoot PD proteome had an overlap of 80% with published PD proteomes and included ∼43% of known high-confidence PD proteins, e.g., the well-established plasmodesmata-located proteins (PDLP) and multiple C2 domains and transmembrane region proteins (MCTP) PD markers (table S3, Figure 2-figure supplement 1 and 2, file S3)(Brault et al., 2019; Thomas et al., 2008). The Arabidopsis PD-enriched fraction included seven Arabidopsis FG-NUPs (Figure 2A). Moreover, inner and outer ring and basket NUPs were found in the high-confidence PD protein list (Figure 2A, C). In total, 20 proteins from different NPC domains were identified. 15 proteins were enriched in PD fractions compared to cell wall and total Arabidopsis protein fractions (Figure 2A, C, Figure 2-figure supplement 1 and 2). Several of the identified nuclear pore components passed the threshold for consideration as high-confidence PD proteins, in particular proteins of the inner and outer rings, core and basket. High-confidence PD scoring was based on enrichment in biochemical PD fractions as well as on an iterative weighting process which factors in features of known PD-localized proteins (Gombos et al. 2023). Separation of PD scores of “likely PD proteins” and likely contaminants (here: plastidic and mitochondrial proteins)(Fernandez-Calvino et al., 2011; Gombos et al., 2023) was significant, allowing to define confidence scores. NUP proteins were also previously identified in PD proteome preparations. In particular NUPs such as FG-NUPs NUP62, NUP214, NUP50a, as well as CPR5, GP210, NUP35, NUP93a, NUP155, NUP205, NUP43, HOS1 (aka HOS1/ELYS), SEC13, and NUP82 were found in proteomics data of cell wall/PD protein fractions from other studies (table S4)(Bayer et al., 2006; Fernandez-Calvino et al., 2011; Gombos et al., 2023; Kirk et al., 2022; Kraner et al., 2017). Biochemical enrichment of PD resulted in enrichment of well-established PD proteins, such as PDLPs and MCTPs (Figure 2-figure supplement 2). As expected, likely contaminants, such as plastidic and mitochondrial proteins were depleted during PD preparation. Very highly abundant proteins, such as cytosolic ribosomal proteins, as well as histones remained high in all fractions. Ribosomal proteins did not show significant differences in abundance between fractions, while histones seemed enriched in cell wall preparations. The abundance of nuclear proteins of the nucleus showed distinct patterns: NUPs were enriched in cell wall and PD fractions compared to total cell extract, as 75% of NUP proteins were more abundant in PD fractions, exceeding the proportions observed in TC (60%) and CW (∼50%) fractions. The abundance of other nuclear envelope proteins was unaffected by the PD purification and showed no enrichment in PD fractions (Figure 2-figure supplement 2). In contrast to NUPs, ER-resident proteins displayed no significant change in abundance in the cell wall fraction compared to total cell extract, yet showed a slight but consistent enrichment in the PD fraction (Figure 2-figure supplement 2). Of note, grouping of the proteins provides an average; individual proteins within each group may deviate from the average pattern. While the detection of a large number of NUPs including FG-NUPs in the PD fractions, even with enrichment, may support a possible role in PD functions, we can not fully exclude that they represent contaminants.

**Figure 2.**
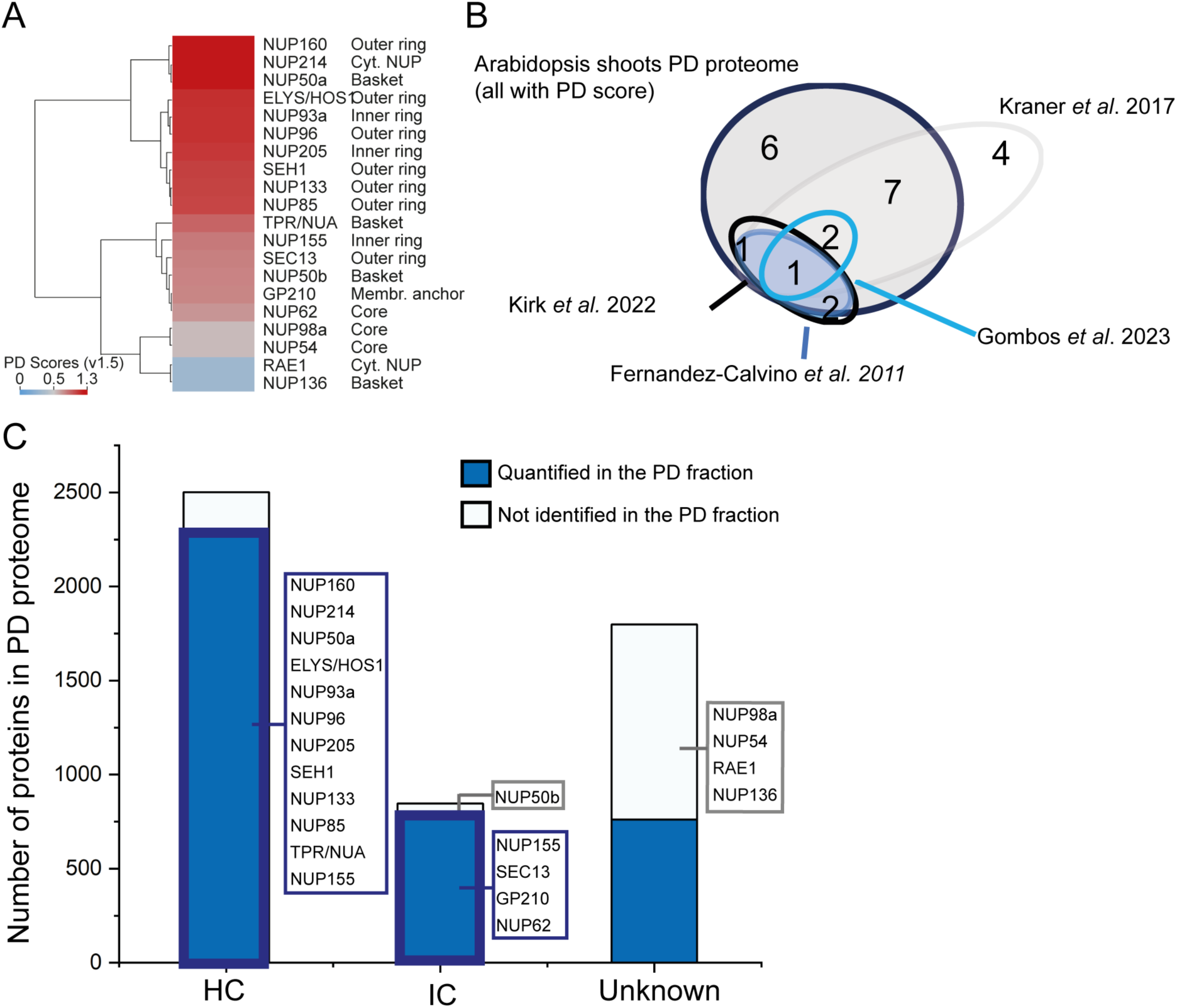
NUPs are found in Arabidopsis shoot proteome and other PD proteomes. **(A),** Hierarchical clustering of PD scores of Arabidopsis NPC components found in PD proteome preparations from 5-week-old Arabidopsis shoot tissues. Clustering based on Euclidean distance of PD scores (v1.5), which were averages of at least three replicate scores from separate tissue preparations. Color scale represents PD score values. PD scores above the high-confidence threshold of 0.5 are indicated in shades of red. (**B**), Comparison of NUPs in Arabidopsis shoot PD proteomes with published PD proteomes. Among the NUPs identified in the Arabidopsis shoot PD proteome, 73.3% (11 out of 15) were also detected in previously published PD proteomes. (**C**), Overview of identified proteins in the PD proteome. High confidence (HC) (PD score > 0.83), intermediate confidence (IC) (0.83 > PD score > 0.63), and Unknown (PD score < 0.63). Purple-edged boxes in HC and MC mark proteins that meet Arabidopsis shoot PD proteome criteria (PD score > 0.63, quantifiable in PD fraction). Blue-edged boxes denote 16 NUPs within the Arabidopsis shoot PD proteome; Gray-edged boxes represent NUPs found also in other proteomes.

### Localization of Arabidopsis FG-NUPs and other NUPs to PD

To explore whether Arabidopsis FG-NUP-mVenus fusions can be detected at PD, candidates were expressed from an β-estradiol-inducible promoter in *N. benthamiana* leaves. Fluorescence derived from FP-fusions of the FG-NUP58, FG-NUP62 and FG-NUP98b overlaid with aniline blue fluorescent puncta (with a significantly higher PD index relative to that for the non-specific cell wall dye propidium iodide, Figure 3A, B, Figure 3-figure supplement 1, table S2), indicating that Arabidopsis FG-NUPs can be targeted to PD. Notably, 12 out of a total of the 16 NUPs tested here localized to puncta in the cell periphery that overlaid with aniline blue fluorescence at pit fields (Figure 3, Figure 4A, B; Figure 3-figure supplement 1, list of 16 tested NUPs: table S2)(Zavaliev and Epel, 2015). Overall, 12 NUPs showed significant enrichment at PD compared to the cell wall marker (Figure 3B, Figure 4B, table S2), including two outer ring NUPs (NUP43, HOS1/ELYS) and two transmembrane NUPs (GP210, CPR5) (Figure 4B, H). Ten NUP fusions for various inner and outer ring and membrane anchor NUPs were also detected in nuclei (Figure 3-figure supplement 1). Fluorescent puncta of aniline blue and FP-fusions of PD protein typically do not provide a perfect colocalization since aniline blue staining does not mark all PD and typically, the overlay is partial since FP-fusions are not detected at all PD even for high confidence PD proteins (Pendle and Benitez-Alfonso, 2015). As an independent line of evidence we observed overlay of the PD marker PDLP5-mScarlet3 and an mCitrine fusion of the putative NUP107-160 complex recruitment factor HOS1/ELYS at PD (Figure 4G). To test for potential artifacts, caused possibly by mistargeting due to overexpression, the concentration of the inducer β-estradiol and the induction time were reduced. In leaves transiently expressing NUP43-mCitrine or CPR5-FP fusions, the fluorescence intensity correlated with the estradiol concentration used, with decreased fluorescence intensity for samples where 2µM estradiol was applied versus the intensity in samples exposed to 20µM estradiol (Figure 4 D,E). Notably, the fluorescence ratio between periphery and nucleus did not differ significantly after expression induction by 2 µM compared to 20 µM β-estradiol (Figure 4F) and PD localization was not eliminated (example for localization of NUP43-mCitrine in Figure 4C; PD index(NUP43, 2µM) = 1.42, PD index(CPR5, 2µM) = 1.40). The resolution of confocal imaging is insufficient to conclude that the NUPs localize inside PD or accumulate in the vicinity. We also cannot exclude that expression of the NUP fusions leads to mis-targeting to PD. Different from data obtained for NUP98a (Gallemí et al., 2016), a C-terminal mVenus fusion to the auto-cleavable NUP98b was fluorescent and localized to PD (Figure 3, Figure 3-figure supplement 1), indicating that either under the condition used in our experiments, the main fraction of NUP98b was unprocessed, or that the C-terminal peptide reassociated with the N-terminal domain of NUP98b, as previously described for the two splice forms of the human NUP98 (Hodel et al., 2002). Out of 16 NUPs tested, four NUPs did not localize at PD, as indicated by non-significant PD indices over the cell wall marker (*p-*values > 0.08, table S2). Out of 35 currently known Arabidopsis NUPs, 19 NUPs were not yet tested here (table S2).

**Figure 3.**
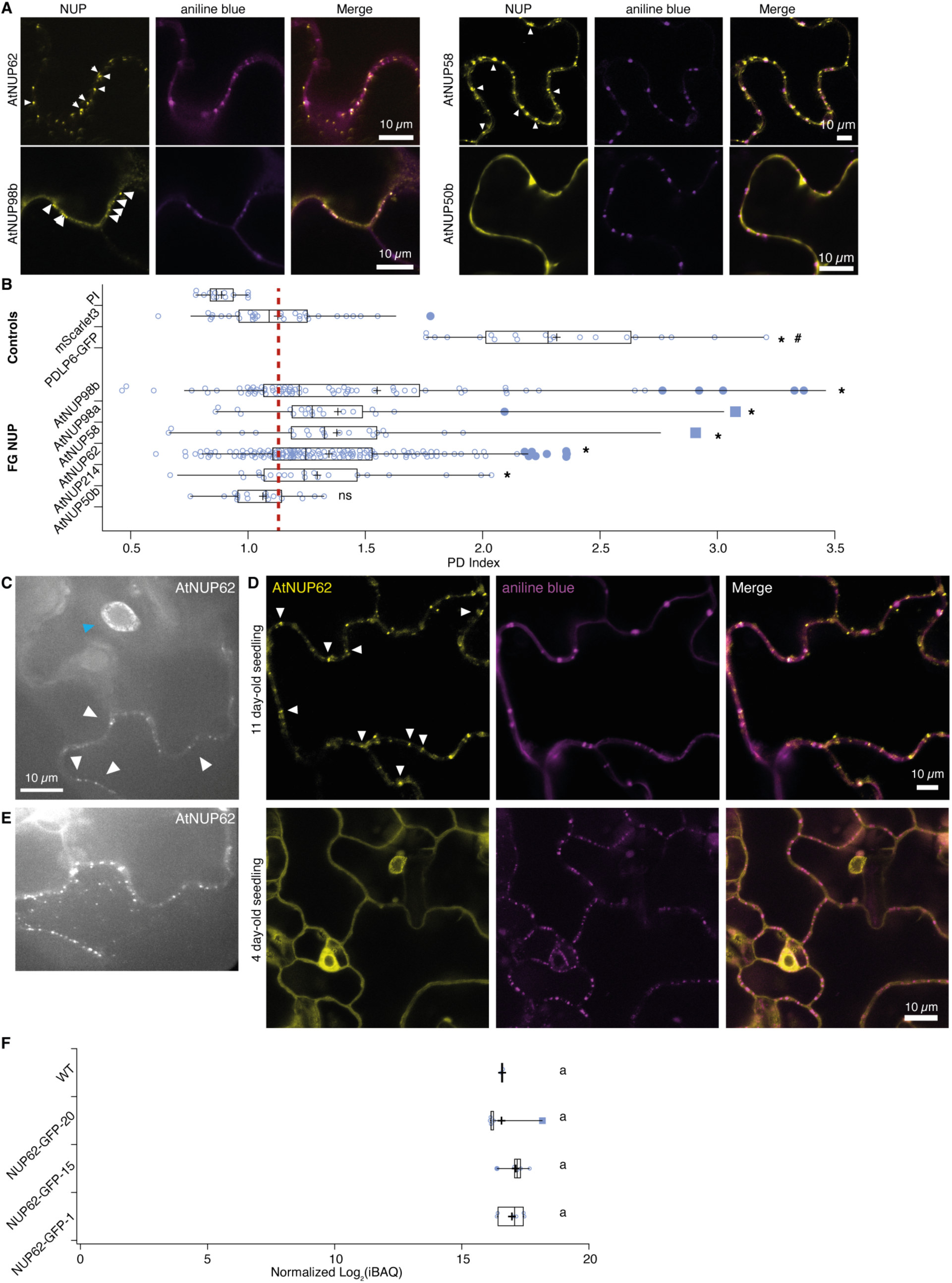
Dual localization of Arabidopsis FG NUPs. **(A)** Localization of selected Arabidopsis FG-NUPs after transient expression under the control of the β-estradiol-inducible XVE promotor in *N. benthamiana* epidermal cells visualized by a single confocal optical section. mVenus was fused to the C-terminus of different NUPs, except for NUP98a for which GFP was fused both N- and C-terminally (GFP-NUP98a-GFP). Aniline blue was infiltrated to detect callose in pit fields as a PD marker. Localization was repeated in three independent experiments with similar results. (**B**) Quantification of the PD localization by PD index. Red dashed line indicates the mean PD index of mScarlet3 (cytoplasmic control). PI, propidium iodide (‘membrane impermeant dye’). Significantly (*p* < 0.05) increased PD indices compared to the cell wall or cytoplasmic controls are indicated by * and #, respectively. Kruskal-Wallis Test *p* < 0.001, *p*-values of Bonferroni-corrected Dunńs test for pairwise comparison are summarized in table S2. Median is represented by vertical line inside the box, mean is represented by the bold **+**. Values between quartiles 1 and 3 are represented by box ranges, and 5^th^ and 95^th^ percentile are represented by error bars. At least 15 images from 3 biological replicates were analyzed for each NUP. Independent localization data are shown in Figure 1-figure supplement 3. (**C-E**) Arabidopsis NUP62-GFP plants expressing NUP62 fused to GFP under its own promoter. (**C**) The localization of NUP62-GFP in 11-day-old cotyledon epidermal cells was visualized by a single confocal optical section (CSU10; 100x oil-immersion objective). Arrowhead: blue nucleus, white peripheral NUP62-GFP. **(D)** Localization of NUP62-GFP in cotyledonar epidermal cells of stably transformed 11-day-old (*top*) and 4-day-old (*bottom*) Arabidopsis lines. Cotyledons were immersed in aniline blue to detect callose in pit fields as PD markers. White arrowheads in D indicate overlay of NUPs and aniline blue. Scale bar: 10 µm. **(E)** Maximum intensity projection of 20 optical cross-sections taken at 1µm intervals of epidermal cotyledon cells expressing NUP62 fused to GFP under its own promoter (11-day-old seedling). Punctate localization of NUP62-GFP can also be found at the epidermis cell surface on the adaxial side of the cell. **(F)** Quantification of NUP62 protein abundance by MS analysis in cotyledons of 14-day old Arabidopsis seedlings. Normalized iBAQ values were used to compare transgenic lines with wild-type (Col-0), n=2-6 technical repeats based on 2-3 independent biological sample preparations. Statistical significance was determined by one-way ANOVA: *p* = 0.5. Different letters indicate statistically significant differences at *p* < 0.05. Mean normalized log_2_ (iBAC) ± SEM: *WT*: 16.58 ± 0.04; *NUP62-GFP-1*: 16.97 ± 0.2; *NUP62-GFP-15*: 17.12 ± 0.2; *NUP62-GFP-*20: 16.56 ± 0.4.

**Figure 4.**
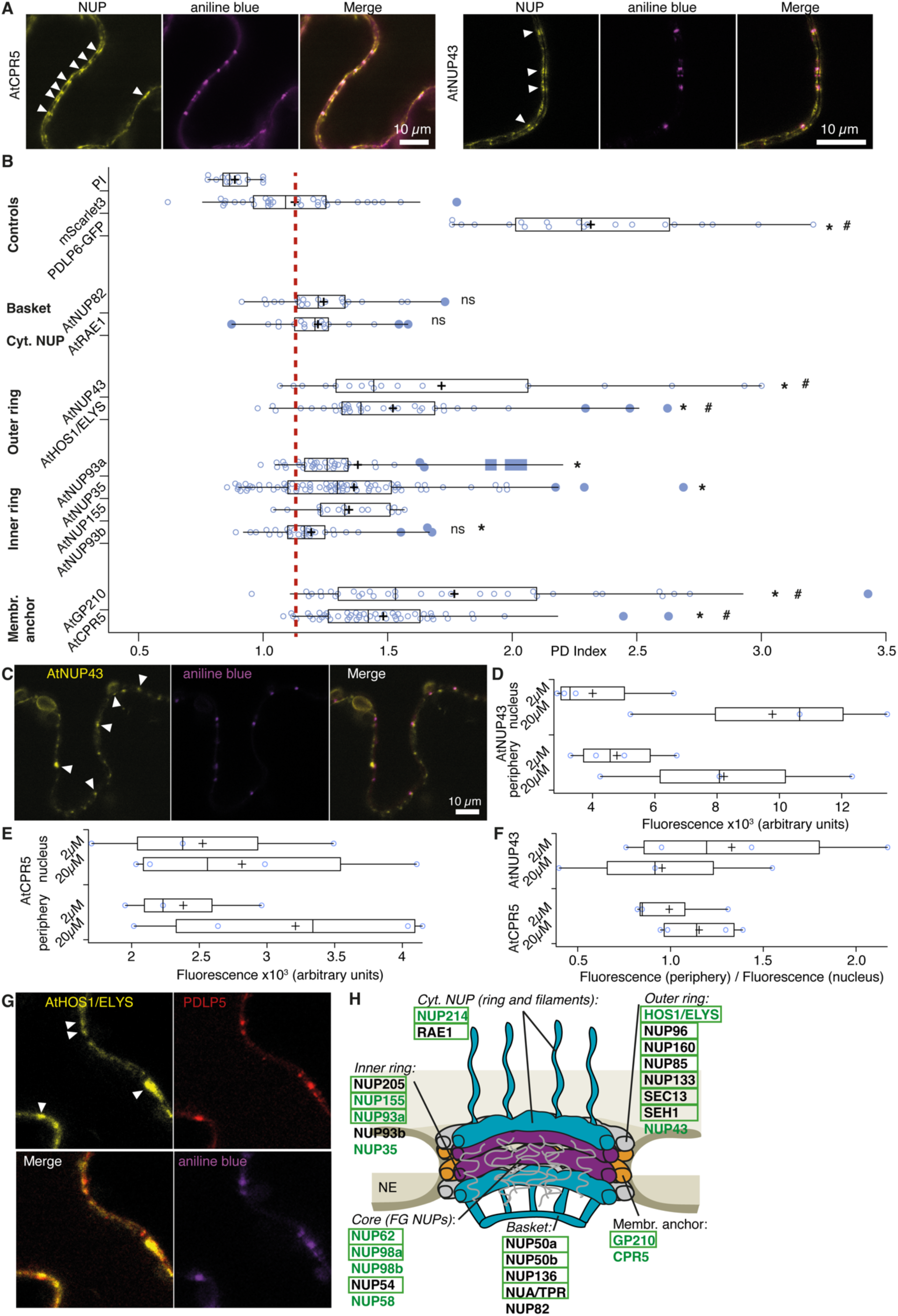
Dual localization of Arabidopsis scaffold and transmembrane NUPs. **(A)** Localization of Arabidopsis transmembrane (CPR5) and scaffold (NUP43) NUPs after transient expression under the control of the β-estradiol-inducible XVE promotor in *N. benthamiana* epidermal cells. mVenus was fused to the C-terminus of different NUPs. Aniline blue was infiltrated to detect callose in pit fields as PD marker. White arrowheads indicate overlay with aniline blue. Localization was repeated in three independent experiments with similar results. (**B**) Quantification of the Arabidopsis NUPs PD localization by PD index. Red dashed line indicates the mean PD index of mScarlet3 (cytoplasmic control). PI, propidium iodide (‘membrane impermeant dye’). Significantly (*p* < 0.05) increased PD indices compared to cell wall or cytoplasmic controls indicated by * and #, respectively. Kruskal-Wallis Test *p* < 0.001, *p*-values of Bonferroni-corrected Dunńs test for pairwise comparison are summarized in table S2. Median: vertical line inside box, mean: **+**. Values between quartiles 1 and 3: box ranges, and 5^th^ and 95^th^ percentile are represented by error bars. At least 15 images from 3 biological replicates were analyzed for each NUP. Independent data in Figure 1-figure supplement 3. Control identical to Figure 2B. **(C**) NUP43-mCitrine localization at PD transiently expressed in *N. benthamiana* epidermal cells at 1/10 β-estradiol concentration and shortened induction time (not exceeding 10h), visualized by a single confocal optical section. White arrowheads: overlay with aniline blue. **(D and E)** Quantification of the fluorescence intensity of NUP43-mCitrine (D) and CPR5-FP fusions (mVenus and mCitrine) (E) in the periphery and the nucleus after induction by 20µM or 2µM β-estradiol. **(F)** Periphery/nucleus fluorescence ratios in *N. benthamiana* epidermal cells transiently expressing NUP43-mCitrine or CPR5-FP fusions after expression is induced by 2 or 20µM β-estradiol. Box plots in (D-F) are based on a smaller subset of confocal images that show nucleus and peripheral fluorescence in one image; for individual conditions: n ≥ 3 images from at least 2 independent repeats. Student’s t-test in (F) 2µM vs. 20µM estradiol: for NUP43, p = 0.26 and for CPR5, p = 0.45. (**G**) Localization of HOS1/ELYS-mCitrine and PDLP5-mScarlet3 co-expressed transiently under the control of the inducible XVE promoter in *N. benthamiana* epidermal cells. Aniline blue was infiltrated to stain callose that accumulates in pit fields. White arrowheads: overlay of HOS1/ELYS-mCitrine and PDLP5-mScarlet3 channels. Localization repeated in three independent experiments with comparable results. All confocal images in Figure3 are single confocal optical sections. (**H)** Scheme of an NPC embedded in the nuclear envelope with all NUPs studied in this work. NUPs listed in B are green boxed. Green letters: NUPs significantly enriched at PD in transient localization assays (Figure 2B, 3B). Green boxed black letters: NUP was identified in the proteome, but was not enriched at PD in transient localization assays. Corresponding AGI locus identifiers in table S2. Cytoplasmic NUP is abbreviated as Cyt. NUP, and membrane anchor as Membr. anchor.

Single copy insertions of GFP-fusions in stable transformants reduce the potential for artifacts caused by overexpression. Analysis of nine independent stably transformed homozygous single insertion Arabidopsis lines expressing a full-length genomic FG-NUP62-GFP fusion under the control of the native NUP62 promoter showed dual localization of the protein to the nuclei and PD. However, PD localization was only detected in mature cotyledons older than 10 days (Figure 3C-E). Notably, the peripheral localization of NUP62-GFP was not restricted solely to PDs, as punctate fluorescence was detected at the cell surface, outside of PD regions (Figure 3E). NUP62 protein abundance in two-week-old cotyledons of the stable NUP62p:NUP62-GFP transformants was not statistically different to NUP62 protein levels in WT (Figure 3F). Notably, the punctate fluorescence at the cell periphery, encompassing both PD-associated and non-PD-associated localization, were not detectable or absent in roots and young leaves of four-day-old seedlings (Figure 3D). However, it cannot be excluded that the GFP fusion impacts NUP62 localization.

Components of the inner ring, NUP93a and NUP155, and the basket, NUP82, were also detected at PD when heterologously overexpressed in *N. benthamiana*, although with lower confidence compared to, e.g., CPR5. Analysis of published data from other groups, who had localized NUPs to nuclei also showed peripheral puncta, which had not been analyzed further (table S5). We assessed whether NUP localization is distinct from ER localization in *N. benthamiana* leaves that heterologously co-expressed NUP43-mVenus and the ER luminal marker mCherry-HDEL. The localization patterns of NUP43-mVenus and of the mCherry-HDEL luminal ER marker were clearly distinct (Figure 5, Figure 5-figure supplement 1). NUP43-mVenus may be associated to the ER, however restricted to subregions of the ER, which partially overlay with aniline blue-labeled pit fields (Figure 5, Figure 5-figure supplement 1). As of now, we cannot exclude that the localization in the periphery is an overexpression artifact due to accumulation in the ER at the orifices of PD since the complexes are too large for translocation (Giordano et al., 2013). Alternatively, the puncta in the periphery could be the consequence of an accumulation, when biotic or abiotic conditions lead to elevated production of NUPs that cannot replace existing NUPs, similar to annulate lamellae (AL). In such conditions, the peripheral accumulation might serve as cellular storage of excessive NUPs. Interestingly, in mammalian cells, depletion of NUP98 led to FG-NUP accumulation sites in the periphery at ER junctions. Such sites of accumulation may serve as NUP storage depots rather than sites of degradation (Pletan et al., 2025).

**Figure 5.**
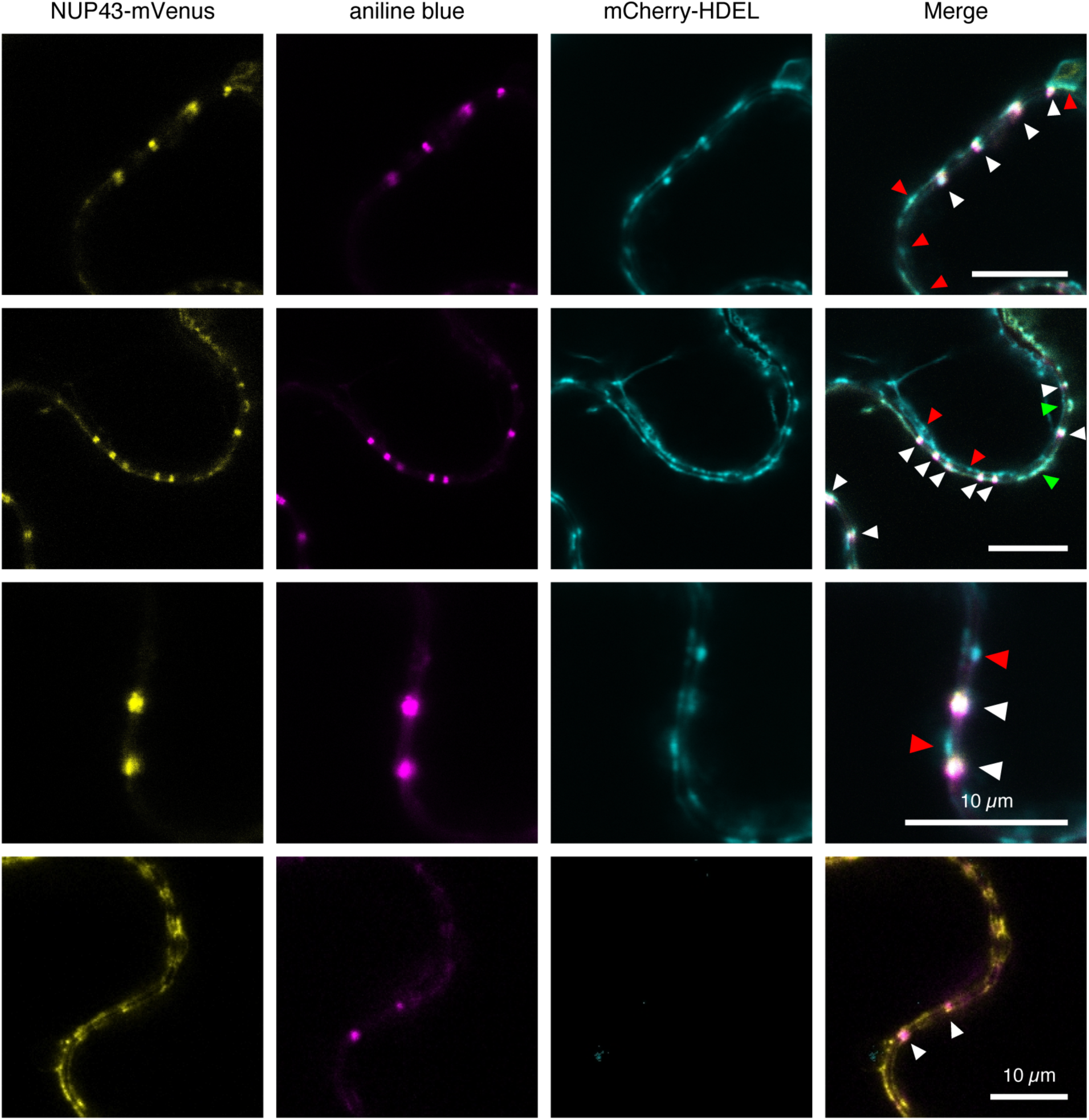
Distinct localization patterns of Arabidopsis NUP43-mVenus and luminal ER marker. Transient co-expression of NUP43-mVenus and mCherry-HDEL in *N. benthamiana* leaf epidermal cells, visualized by a single confocal optical section. Aniline blue was infiltrated to detect callose in pit fields as PD marker. *Bottom,* transient co-expression of NUP43-mVenus without mCherry-HDEL to test for crosstalk. White arrowheads indicate overlay of NUP43-mVenus, mCherry-HDEL. And aniline blue. Red arrows indicate mCherry-HDEL specific localization., without overly with other markers. White arrows (*middle right* panel) indicate overlay of NUP43-mVenus and mCherry-HDEL but absent aniline blue staining. Scale bar indicates 10 µm.

### Topology of CPR5, a potential membrane anchor at both NPC and PD

Since the phase separation domain in the nuclear pore is produced by ten FG-NUPs and PD appear to contain multiple FG-NUPs, it may be necessary to mutate multiple *FG-NUP* genes before it is possible to observe intercellular transport defects (Xiao et al., 2020). It is also conceivable that *knock-out* mutants in multiple *FG-NUPs* may be lethal. We thus rather focused on the green lineage-specific membrane anchor CPR5, a nuclear pore component with a high PD index of 1.48 (Figure 4, table S2). Four potential membrane proteins that anchor the human NPC in the nuclear membrane had been identified: GP210, POM121, NDC1 and TMEM33. Arabidopsis encodes genes for three anchors GP210, NDC1 and PNET1 (homolog of human TMEM209; table S6), while CPR5 had been proposed to serve as an anchor in the equatorial plane of the NPC (Apelt et al., 2016; Gu et al., 2016). Since CPR5 was among the top 5 candidates based on the PD index, one may hypothesize that it may serve as an anchor for an NPC-like structure at PD (Figure 4A, B, Figure 3-figure supplement 1). CPR5, a polytopic membrane protein, is predicted to form a globular helix bundle domain at its N-terminus, flanked by disordered regions and connected to a 105 amino acid α-helix (Figure 6-figure supplement 1). The central helix contains a hydrophobic region, which, together with four shorter hydrophobic helices, forms an approximately 35 Å long transmembrane domain, consistent with the expected thickness of ER membranes. The C-terminus possibly extends into the ER lumen. While it has been considered that CPR5 may play specialized roles, e.g., in defense reactions, it is also conceivable that the phenotypes are secondary defects (Arra et al., 2024; Gu et al., 2016; Ma et al., 2023; Peng et al., 2022), caused by the missing anchoring activity (Apelt et al., 2016; Gu et al., 2016). CPR5 is encoded by a single gene in Arabidopsis and two in rice (Figure 6-figure supplement 2, table S6, table S7)(Arra et al., 2024). In PD, CPR5 could either be embedded into the plasma membrane near or at PD, with the C-terminus facing the apoplasm, or into the ER/desmotubule membrane, with its C-terminus facing the desmotubule lumen (or ER lumen in the vicinity of PD) and the long N-terminal domain pointing into the intermembrane space of PD/cytosol, analogous to the CPR5 topology in the ER-derived nuclear membrane at the NPC. A split-GFP system for topology studies of membrane proteins was used to determine whether the C-terminus of CPR5 faces the luminal side of the ER (Xie et al., 2017). When an N-terminal GFP11 fusion to CPR5 was co-expressed with a free cytosolic GFP1-10, fluorescence was reconstituted, consistent with the N-terminus being located in the cytosol or the cytosolic sleeve of PD (Figure 6A; Figure 6-figure supplement 3). A control using a GFP1-10 domain targeted to and retained in the ER lumen (GFP1-10-HDEL) did not show detectable fluorescence reconstitution (Figure 6B). The complementary fusion of GFP11 to the C-terminus of CPR5 led to reconstitution of fluorescence when co-expressed with an ER lumen targeted GFP1-10-HDEL, but not when co-expressed with a cytosolic GFP1-10 (Figure 6A, B; Figure 6-figure supplement 3). The topological data are consistent with a model in which the C-terminal five-helix-bundle of CPR5 traverses the ER/desmotubule membrane with the C-terminus inside the lumen of the ER/desmotubule, while the extended central α-helix, the predicted unstructured domains, and the smaller helical bundle of CPR5 are located in the intermembrane space between the ER/desmotubule and the plasma membrane (Figure 6-figure supplement 1).

**Figure 6.**
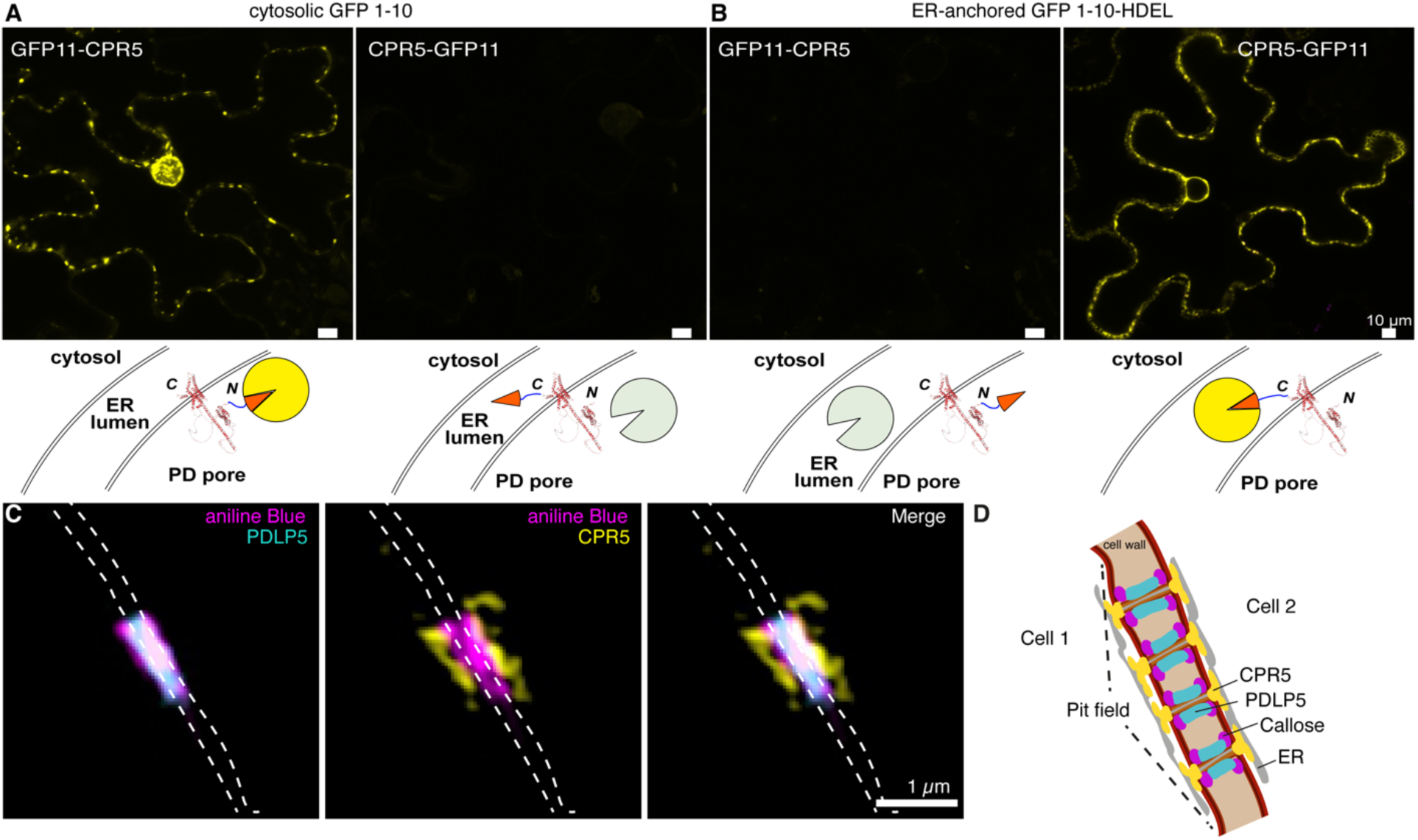
Arabidopsis CPR5 topology and localization of CPR5 near PD orifices. (A and. **B)** Split-GFP membrane topology to test for the orientation of CPR5 in the nuclear pore membrane and the ER at PD by transiently expressing split-GFP fusions in *N. benthamiana,* visualized by single optical sections. (**A)** *(left)* Fluorescence reconstitution at PD between a cytosolic non-fluorescent GFP domain lacking β-sheet 11 (GFP1-10; the circle missing a wedge) and a fusion of the GFP11 to the N-terminus of CPR5 (the red wedge). Since reconstitution occurs at PD, each cartoon indicates reconstitution in the intermembrane space between ER/desmotubule and plasma membrane. *(right)* Cytosolic GFP1-10 and GFP11fusion to C-terminus of CPR5. (**B)** *(left)* ER-retained GFP1-10-HDEL and GFP11 fused to N-terminus of CPR5. *(right)* ER-retained GFP1-10-HDEL and fusion of GFP11 to C-terminus of CPR5. Repeated independently 3 times with comparable results. (**C)** CPR5-mCitrine close to the orifices of PD visualized by a single optical section recorded using Structured Illumination Microscopy in *N. benthamiana* leaves co-infiltrated with XVE:PDLP5-mScarlet3 and XVE:CPR5-mCitrine. Prior to imaging, leaves were infiltrated with aniline blue. PDLP5-mScarlet3: turquoise, CPR5-mCitrine: yellow, aniline blue: magenta. Repeated independently four times with comparable results. **(D)** Cartoon approximately to scale, illustrating PD in pit field between adjacent cells with callose marked magenta, PDLP5 turquoise, CPR5 yellow.

### Localization of CPR5 at the orifices of PD

A hypothetical NPC-like structure could serve as a micropore in the PD channel, or be present close to the PD entrance. To determine CPR5 localization in the PD at higher resolution, we performed structured illumination microscopy (SIM) with a CPR5-mCitrine fusion. CPR5 was found to localize adjacent to aniline blue labeled callose deposition at PD. Therefore, CPR5 may localize to or near the neck region of PD, but not in the central cavity of the PD, where a PDLP5-mScarlet3 was localized (Figure 6C, D; Figure 6-figure supplement 4)(Fitzgibbon et al., 2010). These findings are consistent with a model in which NUPs are present at or close to the PD orifices. We did not observe puncta at both paired neck regions in all cases (Figure 6-figure supplement 4B, top), possibly due to mosaic-like expression in *N. benthamiana*, in which only one of the adjacent cells expressed the fusion protein (model: Figure 6D). CPR5-mCitrine localization was absent at some PD (Figure 6-figure supplement 4B, *bottom*): in 10 images from 3 independent repeats, CPR5-mCitrine and PDLP5-mScarlet3 localized to 76 ± 6% and 81 ± 10% of the aniline blue labeled pit fields, respectively, while CPR5-mCitrine localized to 79 ± 9% of the PDLP5-mScarlet3 puncta (Figure 6-figure supplement 4D). Note that CPR5-mCitrine also localized in the periphery outside PD (Figure 6-figure supplement 4D). To assess whether the CPR5-mCitrine fusion protein is functional in Arabidopsis, we tested whether CPR5p:CPR5-mCitrine (including all introns) expression in the *cpr5-1* mutant background results in a rescue of the severe growth phenotype of the *cpr5-1* loss-of-function mutant (Bowling et al., 1997). Indeed, roots were significantly longer in the two independent transgenic *cpr5-1*/CPR5p:CPR5-mCitrine Arabidopsis lines compared to the *cpr5-1* mutant, and four-week-old plants showed a more WT-like growth phenotype (Figure 7-figure supplement 1, G–I). However, we could not detect fluorescence in 10-14 day old seedlings, which could be due to a variety of reasons, such as cleavage of the FP and degradation of the FP without accumulating elsewhere in the cells. The lack of fluorescence in the transgenic lines requires further investigation.

**Figure 7.**
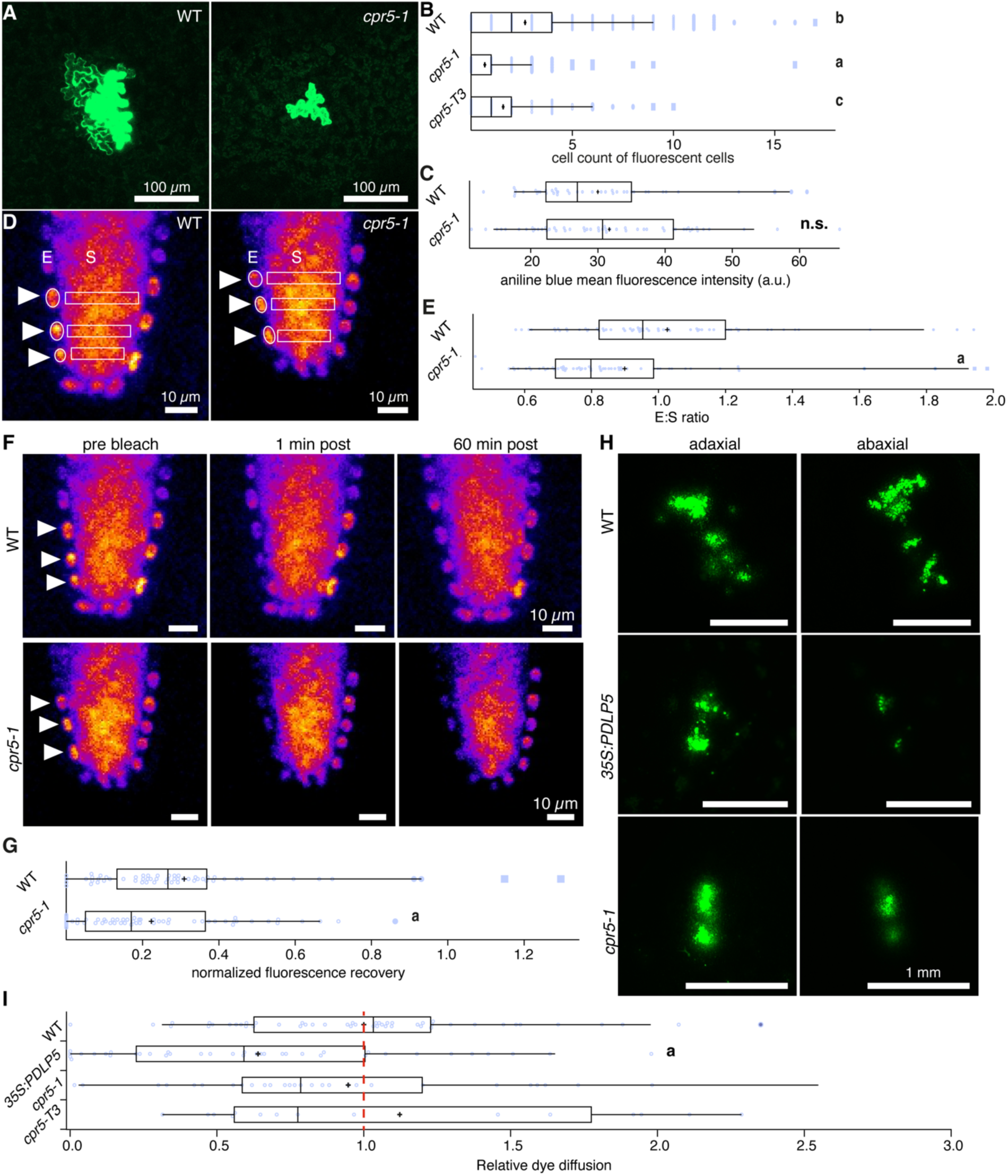
Defects in cell-to-cell transport of kDa-sized cargo in *cpr5* mutants. **(A)** Example maximum intensity projection of 32 optical cross-sections taken at 2.5 µm intervals with intercellular 2xmEGFP movement in wild type (WT, *left*) and *cpr5-1* (*right*) Arabidopsis leaves. Leaves were bombarded with gold particles coated with p35S:mEGFP-mEGFP DNA plasmids 24 h prior to the analysis by confocal microscopy (fusion protein abbreviated as 2xmEGFP). **(B)** Quantification of the mean intercellular spread of mEGFP by counting fluorescent cells in the GFP channel. n_(WT)_ = 594 bombarded cells, 17 independent biological replicates; n_(*cpr5-1*)_ = 278 bombarded cells, 6 independent biological replicates; n_(*cpr5-T3*)_ = 99 bombarded cells, four independent biological replicates. Mean fluorescent cell counts: n_(WT)_ = 2.67, n_(*cpr5-T3*)_ = 1.59, n_(*cpr5-1*)_ = 0.68; median fluorescence cell counts: n_(WT)_ = 2, n_(*cpr5-1*)_ = 0, n_(*cpr5-T3*)_ = 1. Based on Bonferroni-corrected Dunńs test for pairwise comparison after Kruskal-Wallis test: **a** indicates significant difference to WT with *p*_(*cpr5-1*)_ < 10^-15^; **b** indicates significant difference to WT with *p*_(*cpr5-T3*)_ = 0.0004; **c** indicates *p*_(*cpr5-1* vs*. cpr5-T3*)_ = 0.0002. Mean bootstrap analysis with 95% confidence interval (CI) and bootstrap resampling of *B* = 5000: CI_WT vs. *cpr5-1*_ [1 x 10^-5^, 0.001], *p*_(*cpr5-1*)_ = 0002 ; CI_WT vs. *cpr5-T3*_ [1 x 10^-5^, 0.001], *p*_(*cpr5-T3*)_ = 0.0002; CI*_cpr5-1_* _vs. *cpr5-T3*_, *p*_(*cpr5-1* vs*. cpr5-T3*)_ = 0.0002 [1 x 10^-5^, 0.001]. **(C)** Callose quantification. n_(WT)_ = 4 independent experiments, n_(*cpr5-1)*_ = 5 independent experiments, graph summarizes all experiments. Student’s *t*-test *p* = 0.48. **(D)** Sum projections showing SHR–GFP distribution in WT (*left*) and *cpr5-1* mutant (*right*) Arabidopsis root. White circles and boxes indicate positions of endodermis nuclei (E, arrows) and corresponding regions in the stele (S, white boxes). **(E)** Quantification of SHR–GFP endodermis:stele ratio (E:S) for WT and *cpr5-1* mutant. n_(WT)_ = 62 nuclei, n_(*cpr5-1*)_ = 62 nuclei, with 3-4 nuclei analyzed per root. **a** indicates significantly decreased E:S in *cpr5-1* mutant, based on Mann-Whitney U test with *p* = 0.0006. **(F)**, Sum projections showing SHR–GFP distribution in WT (*top*) and *cpr5-1* mutant (*bottom*) Arabidopsis root. **(G)**, Quantification of relative SHR–GFP fluorescence recovery for WT and *cpr5-1* mutant after 1 h. Fluorescence recovery was normalized to a control nucleus in the endodermis (see Methods and Figure 5-figure supplement 1F). n(WT) = 59 bleached nuclei, n(*cpr5-1*) = 59, per root 3-4 nuclei were bleached. Student’s t-test p = 0.037 indicating significantly decreased fluorescence recovery after 60 min. **(H and I)** PD permeabilities examined by CFDA/CFSE-based DANS dye loading assays. **(H)** Epifluorescence reporting on CF diffusion in the epidermis of the adaxial (*left*) and abaxial (*right*) side of the Arabidopsis leaf. WT *(top), 35S*:*PDLP5* (*middle*), and *cpr5-1* (*bottom*). Bars, 1 mm. **(I)** Quantification of CF diffusion in WT and mutant lines. n_(*WT)*_ = 49 plants, 7 independent biological replicates; n_(*35S:PDLP5)*_ = 38 plants, 5 independent biological replicates; n_(*cpr5-1)*_ = 29 plants, 5 independent biological replicates; n_(*cpr5-T3)*_ = 15 plants, 5 independent biological replicates. Fluorescent areas on the abaxial side were identified using auto threshold and Fiji YEN-algorithm with user modifications. The same threshold setting was used for the adaxial side. The extent of dye diffusion was quantified by the ratio between the areal spread of fluorescence on the abaxial side and the areal spread of fluorescence on the adaxial side. **a** indicates significantly different on the basis of *p* = 0.032 level according to Bonferroni-corrected Dunńs test for pair wise comparison following Kruskal-Wallis analysis.

### Intercellular transport defects in *cpr5* mutants

If indeed NUPs form phase-separated barriers at PD to control cell-to-cell transport, and given the defects of *cpr5* mutants in nuclear transport, one may expect that interference with CPR5 function might impair *facilitated and non-facilitated cell-to-cell translocation of specific macromolecules*. Particle bombardment of Arabidopsis leaves was performed to transiently express a tandem-GFP fusion (2xmEGFP; 54 kDa) in two independent *cpr5* loss-of-function mutants (EMS mutant *cpr5-1* (G420D) and T-DNA insertion mutant *cpr5-T3* ; Figure 7-figure supplement 1A, B)(Bowling et al., 1997; Crawford and Zambryski, 2000; Faulkner et al., 2013; Gu et al., 2016; Otero et al., 2016; Sakulkoo et al., 2018; Wang et al., 2017). Cell-to-cell movement of 2xmEGFP was significantly reduced in the *cpr5* mutants relative to wild type (WT) (Figure 7A, B; Figure 7-figure supplement 1A, B). Originally named for its *constitutive pathogen response* phenotype, loss of CPR5 activity can trigger constitutive defense responses that could cause callose deposition and closure of PD (Kohler et al., 2002), however, callose levels were not significantly different (Figure 7C).

The transcription factor SHORT-ROOT (SHR), produced in cells of the stele, migrates into the endodermis by *facilitated cell-to-cell translocation,* followed by nuclear import (Nakajima et al., 2001). Intercellular transport of SHR is crucial for tissue patterning (Koizumi et al., 2012; Nakajima et al., 2001; Winter et al., 2024). To explore whether SHR-GFP (86.6 kDa) movement was impacted in a *cpr5* loss-of-function mutant, we compared the endodermis-to-stele ratio (E:S ratio) of GFP fluorescence intensity in roots of homozygous *shr* mutants expressing pSHR:SHR–GFP in WT, as well as in the *cpr5-1* mutant background. The E:S ratio was significantly reduced in *cpr5-1* compared to WT, consistent with a defect in facilitated SHR transport across PD (Figure 7D, E). Fluorescence recovery after photobleaching (FRAP) of nuclei in the root endodermis of homozygous *shr*/pSHR:SHR lines in *cpr5-1* was significantly lower compared to WT, further supporting that targeted transport across PD is reduced in *cpr5-1* (Figure 7F,G, Figure 7-figure supplement 1F). Notably, no significant difference in SHR–GFP fluorescence intensity was observed in the steles of WT when compared to *cpr5-1* lines (Figure 7-figure supplement 1E). The *cpr5* mutants showed no detectable defect in small molecule transport indicative of preservation of the capability to mediate small molecule transport as shown by ‘Drop-ANd-See’ *trans*-leaf diffusion assays (DANS, carboxyfluorescein: 376 Da; control CaMV*35S:PDLP5*; Figure 7H, I)(Lee et al., 2011). In summary, the intercellular transport assay results indicate that a callose-independent transport mechanism for macromolecules is defective in *cpr5* mutants. While consistent with a possible contribution of NUPs to intercellular transport, we cannot exclude that the reduced transport is due to secondary effects in the *cpr5* mutants, which show rather severe phenotypic defects (Figure 5-figure supplement 1C, D).

## Discussion

Based on the observations that (i) a substantial number of NUPs were identified in PD-enriched cell wall fractions, (ii) 12/35 known plant NUPs, including multiple FG-NUPs, localized to PD, and (iii) intercellular tandem-GFP and SHR transport was reduced in *cpr5* mutants could indicate that components of the NPC may generate a complex that can sustain a similar phase separation domain at PD as found in the NPC, and that phase separation generated by FG-NUPs may play a role in intercellular translocation (Kehlenbach et al., 2023). Previous studies had shown that cargo size cannot be the only criterion for cell-to-cell transport efficiency through PD (Alazem et al., 2025; Campbell et al., 2025; Howell et al., 2024; Kragler et al., 1998; Lee et al., 2003). The evidence for PD localization of NUPs obtained here is intriguing, as it may provide new potential explanations for the underlying mechanism of transport of certain mobile molecules (e.g. RNA-protein complexes) that are larger than the expected size exclusion limit (Tee and Faulkner, 2024). In this context the discovery of a new function of the nuclear transport receptor importin α4 as a PD-associated protein in mediating systemic infection of a plant virus in rice may further support the hypothesis of NPC-like transport functions (Lu et al., 2025).

Nuclear transport experiments using surface modified GFPs and analysis of the properties of nuclear transport receptors indicate that the phase separated domain generated by FG-NUPs in the permeation pathway of the NPC facilitates the transit of cargo that carry arginines, cysteines, histidines, and hydrophobic amino acids at their surface (Ejike et al., 2025; Frey et al., 2018). It will be very informative to test whether similar cargo surface properties are important for the transport through PD. Transport assays combined with pharmacological approaches using compounds such as hexanediol, which disrupt phase-separated domains, and systematic analyses of transport using surface-modified FPs of the same size (Frey et al., 2018; Ng et al., 2023) might help identifying phase separated domains at PD.

In summary we found three independent lines of evidence for the presence of NUPs at or near PD in the context of this work, based on subcellular localization, proteomics, and transport. However, all three independent lines could potentially be the result of artifacts. In particular, we found NUP localization at PD in overexpression conditions and, interestingly, also in stable transgenic Arabidopsis lines with presumably lower protein levels. However, NUP localization in the Arabidopsis lines at PD has only been observed in mature cotyledons and so far not in other tissues such as younger leaves or root. We can not exclude that in the NUP62-GFP lines, PD localization of NUPs is caused by elevated protein levels. Alternatively, it might be possible that localization of NUPs at PD occurs at (natively) elevated protein levels and/or depends on factors such as growth condition, pathogen infection and hormones, which were shown to influence mRNA levels of NUPs (Ascencio-Ibanez et al., 2008; Chen et al., 2012; Narsai et al., 2011; Okuma et al., 2014; Song et al., 2009). Interestingly, dynamic changes in subcellular localization of NUPs in response to environmental stimuli were reported previously, e.g., HOS1-GFP re-localizes from the periphery to the nucleoplasm after cold treatment (Lee et al., 2001). Perhaps the PD localization of NUPs is dynamic, occurring only under certain environmental conditions. To validate the subcellular localization, additional evidence is needed from stable transgenic lines with NUP fusions expressed under the control of the native promoter and terminator, complementing the NUP mutant background, and grown in different environmental conditions. Specific antisera raised against plant NUPs that are suitable for immunolocalization may help to clarify the subcellular localization of NUPs under native conditions. New insights may also be generated by cryo-electron tomography, for which recently suitable protocols have been established (Pöge et al., 2025). An alternative hypothesis that also implicates LLPS for the PD permeability barrier is based on the finding that the linker between the N-terminal C2A and C2B domains (C2A-B linker domain) of multiple C2 domains and transmembrane region proteins (MCTP) found in PD is predicted to be disordered and show strong intrinsic propensity for liquid-liquid phase separation (Dickmanns et al., 2025). Cryo-electron microcopy of plasmodesmata identified assemblies that decorate the desmotubule, which are likely formed by MCTPs, and which could then form an extended permeability barrier in the PD pore (Dickmanns et al., 2025).

There is evidence for NUPs to constitute a permeability barrier between cytoplasm and cilia (ciliary transition zone) in animal cells (Blasius et al., 2019; del Viso et al., 2016; Gu et al., 2016; Takao et al., 2017). However, the exact composition of the proposed ciliary pore complex and its anchoring at the ciliary base remains a matter of debate, as cryo-ET structures of the ciliary base/ transition zone are not yet available (Johnson and Malicki, 2019). Similarly, the assembly of NUPs at PD must differ from that at the NPC because, as described above, desmotubules extend through the central channel of PD and the cytoplasmic sleeve width at PD is smaller than the pore width at the NPC (Ejike et al., 2025; Mosalaganti et al., 2022; Nicolas et al., 2017). The localization of the same NUPs at PD as well as nuclear pores is unexpected, given that one would rather expect gene duplication and diversification to enable independent regulation. Note that not all known NUPs were tested here, and some of those tested did not localize to the PD. The structure, function and regulation of the predicted PD pore complex (PDPC) may differ from that of the NPC.

NUPs seem to also play functional roles outside the NPC in animal cells: FG-NUPs were found in several cyto- and nucleoplasmic membrane-less organelles, such as P-bodies, P-granules, stress granules, and viral-associated organelles. In animals, annulate lamellae, preassembled NPCs in ER stacks, are characteristic for rapidly dividing cells during germ cell development, early embryogenesis and cancerogenic growth and might serve as storage place for NUPs (Kessel, 1992; Penzo and Palancade, 2023). AL have also been found in plant cells (Scheer and Franke, 1972). High NUP levels could result in accumulation within AL. Missing an organellar marker for AL, we cannot exclude that fluorescent puncta derived from NUP-FP fusions in our subcellular localization experiments indicate NUP localization to AL. However, the arrangement of fluorescent puncta did not follow a stacking pattern, which we would expect because of the stacked ER sheets in AL.

PD are challenging research subjects due to the embedding into the cell wall, and the presence of multiple membranes. Based on the similarities between transport properties of PD and NPC, one may hypothesize that they both use similar transport mechanisms. Further characterization will be required to explore mechanisms for delivery of a diverse set of GFP-fused NUPs to PD, and to evaluate their localization and trafficking, e.g. by immunolocalization. Alternatively, accumulation could serve as transient storage of NPC complexes, or are a consequence of overaccumulation at the constriction of ER at PD entrances.

## Materials and Methods

### Plant materials and growth conditions

*Physcomitrium patens* (*P. patens*) ecotype Grandsen cultures were obtained from the *International Moss Stock Center (IMSC)* (Freiburg). *P. patens* was cultured in liquid cultures of modified BCD medium as described (Gombos et al., 2023). The BCD medium consisted of the following solutions which were mixed prior to use: Solution B (0.1 mM MgSO_4_·7H_2_O), Solution C (1.84 mM KH_2_PO_4_), Solution DK (1 M KNO_3_), Solution DF (4.5 mM FeSO_4_·7H_2_O), Trace Element Solution (TES) (614 mg H_3_BO_3_, 55 mg CuSO_4_·5H_2_0, 28 mg KBr, 110 mg Al_2_(SO_4_)3·K_2_SO_4_·24H_2_0, 389 mg MnCl_2_·4H_2_0, 55 mg ZnSO_4_·7H_2_0, 28mg LiCl, 10 mM CoCl_2_·6H_2_0, 28 mg KI and 28 mg SnCl_2_·2H_2_0 in 1 L H_2_0), 500 mM diammonium tartrate and 1 M CaCl_2_. For liquid cultures, 1% (m/v) glucose was added. For propagation of liquid cultures, moss was homogenized and then transferred to 100 ml medium in a 250 ml. Cultures were harvested after two weeks of growth at 21°C under long day conditions (16 h light/8 h dark) at ∼90 µmol m^-2^ sec^−1^, 60% humidity.

Arabidopsis *cpr5-1* (G420D) mutant and *cpr5-T3* (SALK_074631) T-DNA insertion mutant were kindly provided by Dr. Gu (University of California Berkeley) and Professor Chun-Hai Dong (Qingdao Agricultural University), respectively (Bowling et al., 1994; Wang et al., 2017). Arabidopsis Col-0 *shr*/pSHR:SHR–GFP line was described before (Cui et al., 2007) and kindly gifted by Ikram Blilou, KAUST. The pSHR:SHR–GFP/*cpr5-1* in the *shr* mutant background was generated by crossing, and identified in the F2 segregating generation by sequencing of CPR5. For quantification of the SHR-GFP endodermis-to-stele ratio, F5 seedlings of the Arabidopsis lines pSHR:SHR–GFP *cpr5-1* were grown on a glass plate coated with ½ salt strength Murashige and Skoog (MS) media agar [half-strength Murashige and Skoog basal salt mixture (Duchefa Biochemie, Haarlem, The Netherlands), 0.1 % (w/v) 2-(N-morpholino)-ethanesulfonic acid, 1.5% agar, pH 5.6] (Dyachok et al., 2010) for 6 days in short day conditions (10 h light, 14 h darkness), at 21°C with a light intensity to 130 µmol m^-2^ s^−1^. For microparticle bombardment assays and callose quantification, Arabidopsis Col-0, *cpr5-1,* and *cpr5-T3* plants were grown on ½ MS plates in the same conditions as described above for 4 weeks. Nine independent stably transformed homozygous Arabidopsis wildtype lines with single insertions for NUP62-GFP were used for fluorescence imaging (T3 segregants; 3:1 segregation) and grown on ½ MS media supplemented with 1% (w/v) sucrose and 1% (w/v) agar in a growth chamber (16 h light/8 h darkness, 23 °C, 50 µmol m^-2^ s^−1^of white light, 60% humidity). For DANS assay, Arabidopsis WT and T-DNA insertion lines were grown in soil with one plant per pot at 21°C under long day conditions (16 h light/8 h dark) for 3 weeks. For analyses of the Arabidopsis shoot proteome, plants were grown in soil at 21°C under long day conditions (16 h light/8 h dark) at ∼90 µmol m^-2^ sec^−1^, 60% humidity. Respective tissues were harvested after three or five weeks. *Nicotiana benthamiana* (*N. benthamiana*) plants were grown individually in a pot under controlled conditions (16 h light/8 h darkness, 26 °C, ∼90 µmol m^-2^ s^−1^, 60% humidity).

### DNA constructs

For transient expression in *N. benthamiana*, the selected NUP open reading frames (ORFs) were cloned into the binary pAB134-mVenus plasmid (Bleckmann et al., 2010). To generate the ORFs, total RNA was extracted from Arabidopsis leaves using the RNeasy plant mini kit (Qiagen, Hilden, Germany), and DNase I treatment was performed following the manufacturer’s instructions. cDNA was then synthesized using the Maxima H Minus First Strand cDNA synthesis kit (Thermo Fisher Scientific Inc., Waltham, Massachusetts, USA). The ORFs were amplified, incorporating the Gateway attB sites, and purified PCR products were cloned into pDONR221 using a BP clonase reaction (Thermo Fisher Scientific Inc., Waltham, Massachusetts, USA). *E. coli* Stellar cells (Takara Bio) were transformed with the resulting entry vectors. Plasmids with the correct sequence were isolated and then recombined by LR reaction into the binary plant expression vector pAB134-mVenus as C-terminal mVenus fusions under the control of a β-estradiol-inducible promoter (Bleckmann et al., 2010). For localization of PpNUP35.1, Pp3c6_16260V3 was amplified and cloned into the binary pAB134-mVenus plasmid. For SIM experiments, XVE:CPR5-mCitrine and XVE:PDLP5-mScarlet3 were generated using GreenGate (Lampropoulos et al., 2013). CPR5 and PDLP5 genomic sequences were amplified with primers including *Bsa*I restriction sites and ligated to pGGC000 previously digested with *Bsa*I. Resulting vectors were used to generate expression constructs using the GreenGate modules: XVE promoter (module A), CPR5 or PDLP5 (module C), C-mCitrine or mScarlet3 (module D), HSP18.2-Terminator (module E), and the pOLE1:OLE1-RFP seed coat selection marker (Shimada et al., 2010) (module F) were ligated to the linearized binary vector pGGZ003.

Monomeric enhanced GFP (mEGFP) and 2xmEGFP sequences were amplified by PCR from plasmids pENTR-R4R3-mEGFP-C (Addgene #113749) and pGGC025 (Addgene #48830), respectively, using Gateway specific primers adding the attB1 and attB2 sites and cloned into pDONR221 by BP reaction (Thermo Fisher Scientific Inc., Waltham, Massachusetts, USA). The resulting constructs, pDONR221-mEGFP and pDONR221-2xmEGFP were used for LR reaction into the destination vector pGWB502 (Addgene #74844). The p35S:3xmCherry construct was generated using GreenGate by ligating the modules Ca MV 35S promoter (Addgene #48815) (module A), 3xmCherry (Addgene #48831) (module C), HSP18.2-Terminator (module E), and the pOLE1:OLE1-RFP seed coat selection marker (module F) into the linearized binary vector pGGZ003. Topology localization of CPR5 was carried out using the split-GFP system with the vectors kindly provided by Prof. Thordal-Christensen (University of Copenhagen) (Xie et al., 2017). pDONR-CPR5 plasmid was recombined into pDest-GW-GFP11 and pDest-GFP11-GW to generate either N- and C- fusions to the GFP 11th β-sheet using LR Gateway reactions (Thermo Fisher Scientific Inc., Waltham, Massachusetts, USA). The GFP1-10 fragment was co-expressed with either cytosolic localization (GFP1-10) or with ER-luminal localization (GFP1-10-HDEL). Plasmids pEG100-35S:nYFP-CPR5, pEG100-35S:cYFP-CPR5 and pEG100-35S:NUP93a-cYFP used for homo-and hetero-oligomerization were kindly gifted by Prof. Xinnian Dong (Duke University. North Carolina, USA) (Gu et al., 2016). The nYFP fragment contains residues 1-172, and cYFP consists of residues 173-238. A translational fusion NUP62-GFP construct was generated from a genomic clone of *NUP62* (At2g45000), including a 1135-bp region 5’-upstream from the initiation ATG and a 407-bp region 3’-downstream from the stop codon. mEGFP was fused to the NUP62 separated by a GGGGSGGGGSGGGGS-linker (Shimozono and Miyawaki, 2008), just before the stop codon using NEBuilder^®^ HiFi DNA Assembly (NEB). The fluorescent protein-fusion construct was digested by *Not*I and integrated into the pBIN40 binary vector (Miyashima et al., 2011). The final binary vectors were introduced into *Agrobacterium tumefaciens* strain GV3101:pMP90 and used to carry out the transformation of Arabidopsis. We confirmed the nucleotide sequences of all constructs using standard DNA manipulation and sequencing techniques. Primers used in this study are listed in table S8.

### Transient expression in *N. benthamiana* leaves

The *Agrobacterium tumefaciens* strain GV3101 (pMP90, pSOUP), containing the desired construct, was cultured overnight, then diluted to 1/10 and grown until the culture reached an OD_600_ of 0.6-0.8. Cultures were pelleted by centrifuge at 4000 rpm for 10 minutes. Bacteria were subsequently re-suspended in an Infiltration Buffer comprising 0.01 M 2-(N-morpholino) ethanesulfonic acid (MES) at pH 5.6, 0.01 M MgCl_2_, and 150 µM acetosyringone at a final OD_600_ of 0.3. The *Agrobacterium* strain containing the intended construct was then combined with an *Agrobacterium* strain containing the P19 silencing suppressor (OD_600_ 0.1) (Qu and Morris, 2002). The resulting mixture was introduced into the leaves of 4-week-old *N. benthamiana* plants through syringe infiltration. Two days after infiltration, gene expression was induced by spraying infiltrated leaves with β-estradiol solution (20 µM β-estradiol, 0.1% v/v Tween-20). Induction time was dependent on protein maturation times and ranged from 6 to 24 h. To test localization of XVE:NUP43-mCitrine or XVE:CPR5-FP fusions at lower expression levels induction was performed with 2 µM β-estradiol and induction times did not exceed 10 h (2 µM β-estradiol, 0.1% v/v Tween-20).

For split-GFP assays, agrobacterial cultures were suspended in the Infiltration Buffer at a final OD_600_ of 0.3. *Agrobacterium* strains containing constructs split-GFP were co-infiltrated with the P19 silencing suppressor (OD_600_ 0.1) into *N. benthamiana* leaves. For split-GFP assays either GFP11-CPR5 or CPR5-GFP11 and GFP1-10 or GFP1-10-HDEL were co-infiltrated. Samples were imaged 2 days post infiltration.

### Microscopy

#### Subcellular localization of FP fusions by confocal microcopy

Confocal microscopy was performed on a Zeiss LSM 900 (Carl Zeiss, Oberkochen, Germany) equipped with Airyscan GaAsP-PMT detectors and diode lasers using a 40x/1.20 water immersion objective (C-Apochromat 40x/1.20 W Korr FCS). mCitrine or mVenus were excited with a 488 nm laser and emission was collected in a 520-579 nm window. GFP was excited at 488 nm and collected in a 500-550 nm window. mScarlet3 was excited at 561 nm and collected from 579-617 nm (Gadella et al., 2023). To visualize pit fields, *N. benthamiana* leaves expressing a protein of interest were infiltrated with 0.1% (w/v) aniline blue (Sigma Aldrich) solution and then imaged (excitation at 405 nm and 3-5% laser power, emission detected in a 440-480 nm window). Free mScarlet3 was expressed and excited to visualize cytoplasmic localization. Propidium iodide, a cell impermeant dye, stains the apoplasm when short incubation times are used. Thus, for visualization of the cell wall, shortly before the experiment, PI was pipetted on the object slide on the leaf disc (excitation 561 nm, 6% laser power, emission 574-617 nm). Pit fields in cotyledons of 8-day old transgenic NUP62-GFP Arabidopsis seedlings were stained by immersing leaves in 0.5% aniline blue in Sørensen’s phosphate buffer [0.5% aniline blue (w/v) in 200 mM Sørensen’s phosphate buffer prepared with Na_2_HPO_4_ and NaH_2_PO_4_ pH 8, Sigma-Aldrich, Saint Louis, Missouri, USA)]. For confocal imaging of NUP62-GFP Arabidopsis lines, an Eclipse Ti2 inverted microscope (Nikon) equipped with spinning disk confocal unit CSU-10 (Yokogawa) and a sCMOS camera PRIME 95B (Photometrics, Tuscon, AZ, USA) was used. GFP was excited with a 488-nm laser, and emission was collected with a 525/50-nm filter (Semrock, Rochester, New York). Images were taken with a 100x oil-immersion lens (CFI Plan Apo Lambda 1.45 NA, Nikon).

#### Structured Illumination Microscopy

For structured illumination microscopy (SIM), only pit fields that were not deeply embedded in the plant tissue and most proximal to the cover glass close to the objective, were observed. To do so, a *N. benthamiana* leaf disk (2-mm diameter) was immersed in water and pressed between an object slide and a high-precision coverslip (1.5H, 18 x 18 mm; A. Hartenstein) using a magnet. The sample chamber was sealed with a flexible mounting adhesive (Fixo gum; Marabu). Z-stacks with a scaling of 149-nm steps were obtained using an Elyra PS1 (Zeiss Microscopy GmbH, Oberkochen, Germany) equipped with a C-

Apochromat 63x/1.2 water-immersion M27 objective. Frame size was 512 x 512 pixels. PDLP5-mScarlet3, CPR5-mCitrine, and aniline blue were excited with 561 nm, 488 nm, and 405 nm laser wavelengths, respectively. Detection filters were set for the different channels: aniline blue: bandpass (BP) 420–480/LP 750; CPR5-mCitrine: BP 495–570/LP 750; PDLP5-mScarlet3: BP 570-650/LP 655. In initial control experiments with the Elyra PS1, we observed more puncta in the green channel after excitation at 488 nm, after having used the 405 nm laser to excite aniline blue. These additional puncta in the green channel precisely overlayed with the puncta observed in the aniline blue channel. The identification of fluorescent puncta in this channel in response to excitation of aniline blue might be the result of photoconversion to longer wavelengths. To exclude possible artifacts due to photoconversion of the aniline blue, which could lead to emission in the channels used for detection of mCitrine, sequential excitation was performed at 561, 488 and 405 nm. The grating periods (grid size) for structured illumination were set to 34, 34, and 28 µm, respectively, for the three different laser wavelengths. The number of grid rotations was set to 3 rotations. Camera exposure time ranged from 49-128 ms. The setup was operated using Zeiss ZEN 2.3 SP1 software. An internal deconvolution program for SIM z-stacks was used in 3D mode with maximum isotropy, zero sectioning, and the baseline shift active for all three excitation channels. The noise filter was initially run in automatic mode, but was set to a unit below the automatic setting for the actual processing.

#### Callose quantification in Arabidopsis seedlings

CPR5 is involved in the activation of pattern triggered immunity (PTI) and in the repression of effector-triggered immunity (ETI) (Ma et al., 2023; Wang et al., 2014). The *cpr5-1* mutant had previously been reported to show constitutive resistance to two virulent pathogens (Bowling et al., 1997). An important plant defense mechanism against pathogens is induction of callose deposition at PD. Elevated callose deposition in the *cpr5-*1 mutant after wounding has been reported (Kohler et al., 2002). To test whether callose deposition was increased relative to WT under the growth and experimental conditions used here, PD callose levels in WT and *cpr5-1* Arabidopsis seedlings were quantified. Leaf 5 of young seedlings (8-12 leaf stage) was stained to minimize variability. Leaves were detached with a forceps, immersed in 1% aniline blue diluted in water [1% aniline blue (w/v); Sigma-Aldrich, Saint Louis, Missouri, USA] and infiltrated by applying a vacuum in a 20 ml syringe. Aniline blue fluorescence was observed on a Zeiss LSM 900 (Zeiss Microscopy GmbH, Oberkochen, Germany) C-Apochromat 40x/1.20 W Korr FCS and GaAsP-Pmt1 detector with excitation at 405 nm, 5% laser power and gain at 632 V. Emission was detected from 400-500 nm. Imaging parameters were consistent across all experiments. Per experiment, 3-4 leaves were assayed per genotype, 4 images were acquired per leaf. Four independent experiments were performed for WT and 5 for *cpr5-1*. Mean aniline blue fluorescence intensity was quantified with Fiji using the calloseQuant macro (calloseQuant settings: Method A, peak prominence = 55 and measurement radius = 6) (Huang et al., 2022). Regions of Interest (ROIs) assigned to non-PD regions by the Fiji macro were removed manually. Mean aniline blue fluorescence per image was plotted.

### Plasmodesmatal proteome

#### Preparation of *A. thaliana* plasmodesmatal (PD), cell wall (CW) and total cellular proteins (total cell extract, TC) fractions

Various Arabidopsis shoot tissues, including cauline leaves, meristem, and rosette leaves, were collected individually. Each sample pool consisted of at least six plants. PD preparation involved a minimum of three pools of the same tissue type. Tissues were freshly excised and promptly frozen in liquid nitrogen. The PD preparation protocol was based on two previously published methods (Faulkner and Bayer, 2017; Kraner et al., 2017). Each pool of samples was ground in liquid nitrogen and incubated in PD extraction buffer [150 mM NaCl, 10 mM EDTA, 1 mM DTT, 1% (v/v) Triton X-100, 0.5% (v/v) sodium deoxycholate, 0.1% (w/v) SDS, 10% (v/v) glycerol, and 100 mM Tris-HCl, pH 8] with 0.5% Protease Inhibitor Cocktail (PIC, P9599, Sigma-Aldrich), 5 mM DTT, 1 mM PMSF, 50 mM NaF, 1 mM Na_3_VO_4_, 1 mM benzamidin and 4 µM leupeptin. For the cell wall fractions, two rounds of grinding in liquid nitrogen efficiently removed soluble protein contaminants and better prepared cell walls for digestion. The pellet was washed at least four times with a low concentration of detergents [150 mM NaCl, 10 mM EDTA, 0.1% (v/v) Triton X-100, 10% (v/v) glycerol, and 100 mM Tris–HCl, pH 8]. After the last washing step, the sample was centrifugated at 1500 x g at 4 °C with swing-out bucket (Sigma 3-30KS, Rotor 11390 & 13150). Cell wall fractions (CW; 100-200 µl) were collected from the final pellet. The rest of the CW pellet was digested in cell wall digestion buffer (1.4% cellulase, mannitol 300mM, 10 mM CaCl_2_, 10 mM KCl) with 1 mM DTT and 0.5% PIC (P9599, Sigma-Aldrich) at room temperature for 1 h. After centrifugation, the supernatant was collected and ultracentrifuged for 80 min at 4 °C and 110,000 x g (Beckman XPN-80, Rotor SW41Ti). After ultracentrifugation, the resulting pellet was washed twice with TBS buffer (2.5 mM KCl, 140 mM NaCl and 20 mM Tris-HCl, pH 7.4 with 0.5% PIC). The final pellet was PD fraction (PD). Total protein (total cell extract, TC) was extracted according to He *et al*. (He et al., 2022). All fractions were dissolved in 8M UTU (6 M urea, 2 M thiourea, Tris-HCl pH 8.0) before in-solution tryptic digestion. For Arabidopsis cotyledon protein extraction and purification, cotyledons from wild-type (Col-0) and NUP62-GFP single insertion transgenic lines were harvested and immediately transferred to 1.5 ml microcentrifuge tubes. Samples were prepared in three biological replicates with two technical replicates each. Tissue was homogenized in RIPA buffer and centrifuged at 14,000 × g for 15 min at 4°C. The resulting supernatant was carefully transferred to a fresh tube. For protein precipitation, 1/20 volume of 2.5 M sodium acetate (NaOAc, pH 5.0) was added, followed by 5 volumes of ice-cold ethanol (EtOH). Samples were mixed by gentle inversion and incubated at room temperature (24°C) for 3 hr, then centrifuged at 21,000 × g for 20 min. The protein pellet was air-dried briefly and subsequently resolubilized in UTU buffer (6 M urea, 2 M thiourea).

#### Protein clean-up, trypsin digestion and peptide desalting

A single-pot, solid-phase-enhanced, sample-preparation (SP3) technology (Hughes et al., 2019) with trypsin digestion was used to remove the detergent in the protein extracts. Disulfide bridges were reduced by an excess of DTT (6.5 mM), and alkylation of free cysteines was done by iodoacetamide (27 mM). SpeedBead Magnetic Carboxylate Modified Particles (SP3 beads, 45152105050250 / 65152105050250, GE Healthcare) were pre-washed by Milli-Q water. SP3 beads (50 µg/µl) were added to the protein sample in a ratio of 1:10. 50% ethanol was used for inducing protein-bead binding. After incubating 15 min at 24 °C, 1,000 rpm constant shaking, the beads were washed three times with 80% ethanol and subjected to trypsin digestion [100 mM Ammonium bicarbonate, trypsin (trypsin: protein ratio 1:100, V5113, Promega)] for 12-14 h at 37 °C. Digested proteins were acidified to pH 2 using TFA and were then desalted over C18 STAGE tips (Rappsilber et al., 2003).

#### LC-MS/MS analysis of peptides

Peptide mixtures were analyzed by HPLC system nanoflow Easy-nLC (Thermo Scientific, Waltham, Massachusetts) and Orbitrap hybrid mass spectrometer (Q-exactive, Thermo Scientific). Peptides were eluted from a 75 µm x 25 cm analytical C18 column (PepMan, Thermo Scientific) on a linear gradient running from 5% to 90% acetonitrile over 70 min. Proteins were identified based on the information-dependent acquisition of fragmentation spectra of multiple charged peptides. Up to ten data-dependent MS/MS spectra were acquired in the linear ion trap for each full-scan spectrum acquired at 60,000 full-width half-maximum (FWHM) resolution.

#### Peptide and protein identification

MaxQuant version 2.2.0.0 was used for raw file peak extraction and protein identification against the Arabidopsis TAIR11 database (35,386 entries). The following parameters were applied: trypsin as cleaving enzyme; minimum peptide length of six amino acids; maximal two missed cleavages; carbamidomethylation of cysteine as a fixed modification; N-terminal protein acetylation, and oxidation of methionine as variable modifications. Peptide mass tolerance was set to 20 ppm, and 0.5 Da was used as the MS/MS tolerance. Multiplicity was set to 1. For label-free quantification, retention time matching between runs was chosen within a time window of two minutes. iBAQ quantification was selected within label-free quantification. The peptide false discovery rate (FDR) and the protein FDR were set to 0.01, while site FDR was set to 0.05. Hits to contaminants (e.g. keratins) and reverse hits identified by MaxQuant were excluded from further analysis. The mass spectrometry proteomics data have been deposited to the ProteomeXchange Consortium via the PRIDE partner repository (Deutsch et al., 2020) with the dataset identifier PXD049466.

#### PD score (V1.5)

The PD scoring (v1.5) was derived from the earlier PD score equation by Gombos *et al*. (Gombos et al., 2023) with the modification that its components enrichment score and feature score were scaled between 0 and 1. The proteingroups.txt output from Maxquant was analyzed using Perseus. Protein iBAQ values were normalized using z-scores, and the values for PD, CW, and TC fractions were averaged across tissues (file S3). The ratios PD/CW and PD/TC were then calculated based on the average values obtained for each fraction. The PD enrichment score for each protein is the sum of PD/CW and PD/TC. The score was then scaled to a range of 0 to 1, based on the proportion of the distance to the minimum score. Regardless of the enrichment score, the feature score relies on the frequency of protein functions, subcellular localizations and PFAM domains (available at https://pfam.xfam.org/) identified within the high-confidence dataset and validated PD candidates (Gombos et al., 2023; Johnston et al., 2023; Kirk et al., 2022). Features of known PD proteins were extracted and weighted against the entire Arabidopsis proteome. Structural features included Pfam domains PF00722 (GHL), PF06955 (XET_C), PF08372 (PRT_C), PF00335 (Tetraspanin), and PF00168 (C2 domain). Subcellular localization features included plasma membrane (PM), endoplasmic reticulum (ER), extracellular space (EX), and cell wall (CW). Functional features were assigned according to MapMan categories bin 10, 15, 26, and 30. Similar to the enrichment score, the feature score was also normalized to a scale from 0 to 1, based on the proportion of the distance to the minimum score across all *Arabidopsis* proteins (30843 entries). For each protein, the PD score (v 1.5) is obtained by summing the transformed PD enrichment score and the feature score, resulting in a range from 0 to 2. A higher PD score (1.5) indicates a greater likelihood that the protein is a PD protein.

### Intercellular Transport assays

#### Microparticle bombardment

To test passive diffusion of 2xEGFP, microparticle bombardment was performed to deliver DNA coated gold microcarriers into single epidermal cells. 60 mg of 1.0 µm gold microcarriers (Bio-Rad Laboratories, Hercules, CA, USA) were added into a 1.5-mL microcentrifuge tube and washed with 70% ethanol before use. After the final washing step, 1 mL of 50% glycerol was added, and the tube was sealed using Parafilm^®^ (Amcor). The suspended gold particles (35 µL) were labelled with 2 – 2.5 µg of plasmid DNA (pGWB502-p35S:2xmEGFP) in the presence of 0.1 M spermidine and 2.5 M CaCl_2_. Tubes were vortexed for 1 min and put on ice for 1 min. Coated gold particles were centrifuged at 1,500 x g for 30 sec. The particles were washed twice with 70% ethanol. After the second washing step, 35 µL of 100% ethanol was added and vortexed until the pellet completely dispersed. This gold suspension (10 µL) was transferred onto a macrocarrier (Bio-Rad Laboratories, Hercules, CA, USA). Bombardment was performed using the PDS-1000/He Biolistic Particle Delivery System (Bio-Rad, #1652257) as described (Tee et al., 2022). To visualize bombardments, 2xmEGFP was excited with a 488-nm laser and emission detected at 495-548 nm, while 3xmCherry was excited with a 561-nm laser and emission detected at 548-617 nm. Cells showing fluorescence in the GFP channel were counted manually on maximum projections. To identify the initially bombarded cell, Arabidopsis leaves were bombarded with gold particles coated with pGWB502-p35S:3xmCherry (84 kDa, unable to move passively through non-specific trafficking) and pGWB502-p35S:2xmEGFP. Fluorescence intensity profile plots across transects were generated using Fiji software package (Schindelin et al., 2012) to analyze the correlation of the fluorescence intensity of mCherry (bombarded cell) and mEGFP (Figure 7-figure supplement 1).

#### SHR-GFP endodermis-to-stele ratio and Fluorescence Recovery After Photobleaching (FRAP) analysis

SHR–GFP was detected in root endodermis and stele using a Zeiss LSM 900 equipped with Airyscan GaAsP-PMT detectors and diode lasers using a C-Apochromat 40x/1.20 W Korr FCS water objective. For quantification of SHR-GFP, a brief Z-stack series of up to 60 sections of short intervals (2.1 μm) around the ROI were captured, and the Sum-Projection of the volume was analyzed using ImageJ. The background fluorescence was subtracted from the average pixel intensity in the circled region in the endodermis (E) and from the average pixel intensity of the corresponding boxed region in the stele (S) (see Figure 7D). The ratio between endodermis and corresponding stele region was calculated (E:S). Multiple samples were taken for each root to determine the average ratio.

The SHR–GFP fluorescence was bleached in endodermal cells using the 488 nm laser at full laser power for 150 iterations, which lead to around 50-70% reduction in fluorescence. 1-3 cells in the same cell layer were bleached at the same time for each root. To acquire recovery images, laser power was set to 2% to avoid bleaching. The recovery images of the bleached areas were captured every 20 min to reduce the possibility of additional bleaching from frequent scanning. To ensure the identification of the same region of interest during the time-course imaging, a brief Z-stack series of up to 60 sections with short interval (2.1μm) around the region of interest were captured. In this way, the same region should be included in the series of images and can be re-tracked later for analysis. For each time point the Sum-Projection of the same volume was analyzed. The background subtracted fluorescence intensity ratio (endodermis bleached nucleus/endodermis control nucleus) before bleach, after bleach and after 60 minutes was determined using ImageJ (see also Figure 7-figure supplement 1F for positions of ROIs). Time 0 post-bleach fluorescence was set to 0 and all measurements were normalized to this point by subtracting the time 0 post-bleach fluorescence signal from all of the pre-bleach and the recovery values. The percent recovery was calculated using the normalized values. For analysis of intracellular FRAP, a frame size of 512 × 512 pixels with a scanning speed of 0.93 ms per frame (1 Hz) and bi-directional scanning was used to accelerate the scanning rate for each frame.

#### DANS assay

For Drop-ANd-See (DANS) assays, 1 μl of 10 mM 5-(and 6)-carboxy-fluoresceindiacetat-acetoxymethylester (CFDA) in 0.002 % Silwet® was pipetted on the adaxial side of a rosette leaf and removed after 5 min. The 4^th^ and 24 h later also the 5^th^ rosette leaf was used. For microscopy, the leaf was cut off, washed in water, and mounted between two cover slips. Fluorescence was observed under an Axio Zoom.V16 (Zeiss Microscopy GmbH, Oberkochen, Germany) equipped with a X-Cite Xylts (Excelitas Technologies, PA, USA) LED light source, using a GFP broad bandpass filter cube (excitation: 470/40 nm, beam splitter: FT495 nm, emission: 525/50 nm). Images of the adaxial- and abaxial side were taken using Orca-fusionBT digital camera (C15440, Hamamatsu, Japan). Imaging parameters were consistent across experiments. To identify the fluorescent area, thresholding was performed in ImageJ with the same threshold value for adaxial-and abaxial sides. The diffusion is quantified by ratio between the area above the intensity threshold on the abaxial side and the adaxial side. Experiments of the same day were normalized to the WT experiments of the same day.

### Bioinformatics

#### FG repeat analysis

To identify FG repeat proteins, *A. thaliana* and *P. patens* peptides encoded by gene models were downloaded from Araport 11 and Phypa_V3 genomes and screened, respectively. FG occurrences were counted in a sliding window of 200 amino acids (aa), and the maximum number of FG repeats per window was determined using a custom python script (Supplementary Script 1 and 2). Within the same script, the number of GLFG and xFxFG motifs in each peptide were counted (table S1).

#### PD index quantification

Overlay of fluorescence from NUP-FPs and aniline blue was quantitatively scored using the PD index (Grison et al., 2019). To minimize bias in quantification, we employed our recently developed PD index script (Gombos et al., 2023). Briefly, clear PD sites were cropped, and channels were separated. PD were identified using auto threshold and Fiji YEN-algorithm, with user modifications. Center coordinates of PD were determined, and square ROIs were set. The PD index was calculated by dividing mean intensity values of PD ROIs by background ROI values. Background ROIs were defined as areas of weak aniline blue fluorescence. Regions previously recognized as PD were automatically omitted from the positioning of background ROIs. Image processing was used for background ROI placement. The software Igor Pro (version 9) was used for data plotting and SPSS was used for statistical analyses.

#### Structure prediction for CPR5

The structure of CPR5 was predicted using Alphafold monomer version 2.3.1 (Jumper et al., 2021).

### Phylogenetic analyses

The tree shown in Figure 6-figure supplement 2 was generated from 68 amino acid sequences as listed in table S7. Full-length sequences were imported into the Molecular Evolutionary Genetics Analysis (MEGA) package version 11, and aligned by CLUSTALW (Tamura et al., 2021). All positions with at least 85% site coverage were used. Phylogenetic analyses were conducted using the Maximum Likelihood method. Based on the suggestions of the “find best protein model (ML)” tool in MEGA 11, the lower BIC (Bayesian Information Criterion) model was selected, which was the JTT matrix-based model (Jones et al., 1992). A discrete Gamma distribution was used to model evolutionary rate differences among sites (five categories, +G, parameter = 6.413). Initial trees for the heuristic search were obtained automatically by applying Neighbor-Joining and BioNJ algorithms to a matrix of pairwise distances estimated using the JTT model, and then selecting the topology with a superior log likelihood value. The bootstrap consensus tree inferred from 1000 replicates was taken to represent the evolutionary history of the analyzed genes. Branches corresponding to partitions reproduced in less than 50% bootstrap replicates were collapsed.

#### Statistics

In the PD proteome from Arabidopsis leaf tissue, we identified 5145 proteins from PD, CW, and TC fractions, with 3347 proteins having a PD score > 0.63. Among them, 3062 proteins were quantifiable in the PD fraction, constituting our Arabidopsis shoot PD proteome. Among these, 2268 proteins were categorized as high-confidence PD proteins, while 792 proteins fall into the medium-confidence category (file S3). The PD index is based on the analysis of at least 15 images from three independent repeats for each NUP and negative control. The quantification of the intercellular spread of 2xmEGFP by counting fluorescent cells in the mEGFP channel is based on the analysis of n_(WT)_ = 17 independent biological experiments with in total 594 bombarded cells; n_(*cpr5-1*)_ = 6 independent biological replicates with in total 278 bombarded cells; n_(*cpr5-T3*)_ = 4 independent biological replicates with in total 99 bombarded cells. Kruskal-Wallis test with Bonferroni-corrected Dunńs test for pairwise comparison was used to test for significant differences in mean cell counts between WT, *cpr5-1*, and *cpr5-T3* with *p*_(WT vs. *cpr5-1*)_ < 1×10^-10^, *p*_(WT vs. *cpr5-T3*)_ = 0.025, and *p*_(*cpr5-1* vs*. cpr5-T3)*_ *=* 0.00005. The *p-*values are summarized in table S2. The callose quantification is based on n_(WT)_ = 4 independent experiments, n_(*cpr5-1)*_ = 5 independent experiments. Student’s *t*-test indicated a non-significant difference with *p* = 0.48.

SHR-GFP endodermis-to-stele ratio is based on the quantification of n_(WT)_ = 62 and n_(*cpr5-1*)_ = 62 different endodermis nuclei and corresponding stele regions, while per root 3-4 different nuclei and corresponding stele regions were quantified. For endodermis-to-stele ratio, Mann-Whitney U test indicated a significant difference between WT and *cpr5-1* background with *p* = 0.0006. For baseline stele fluorescence, Mann-Whitney U test indicated no significant differences between s*hr* and s*hr x cpr5-1* with *p* = 0.074. The quantification of SHR-GFP fluorescence recovery was normalized to a control nucleus in the endodermis (see Methods and Figure 7-figure supplement 1F) and is based on n(WT) = 59 and n(*cpr5-1*) = 59 bleached nuclei, while per root 3-4 nuclei were bleached. Student’s T-Test indicated a significant difference between WT and *cpr5-1* background with p = 0.037.

For the DANS data, Kruskal-Wallis test with Bonferroni-corrected Dunńs test for pairwise comparison was used to test for significant differences between WT and mutant lines. All statistical test were performed with SPSS. In all box plots median is represented by vertical line inside the box, mean is represented by the bold **+**. Values between quartiles 1 and 3 are represented by box ranges, and 5^th^ and 95^th^ percentile are represented by error bars. Data was plotted using Igor Pro (version 9).

## Supporting information

Supp Script 2

Supp Script 1

## Acknowledgments

We would like to dedicate this manuscript to the ‘champion of plasmodesmata’, William J. (Bill) Lucas (UC Davis). We thank Madlen Somsich (HHU) for project coordination and management. This work has received funding from the European Research Council (ERC) and the Marie Skłodowska-Curie Actions under the European Union’s Horizon 2020 research and innovation program (Grant agreements ‘SymPore’ No. 951292 to WBF and “PDgate” No. 101023981 to MM, respectively). ARE was supported in part by funds from the Interdisciplinary Graduate and Research Academy (iGRAD) Düsseldorf (Deutsche Forschungsgemeinschaft (DFG, German Research Foundation), DFG, project ID 391465903/GRK 2466). Research was also supported by the Deutsche Forschungsgemeinschaft (DFG, German Research Foundation) under Germanýs Excellence Strategy – EXC-2048/1 – project ID 390686111, and the Alexander von Humboldt Professorship, to WBF.

## Data availability

Source data have been assigned a DOI (http://doi.org/10.60534/46qra-6bw98) and are accessible via https://git.nfdi4plants.org/projects/2188. Materials will be made available under MTA; some materials were obtained from others as marked and require permission from the respective authors.

## Contributions

WBF conceived of the study. J.E. performed the bioinformatic analyses; L.X., S.G. and W.S. performed proteomics and data analysis; M.D. analyzed the structure and generated Figure 2-figure supplement 1; S.H. helped to establish and advised on SIM. T.M.S, M.P., M.M., J.O.E., N.P., M.N., G.V.D, C.G. and A.D. performed imaging and/or transport studies; R.S. advised on imaging and data analyses; M.M. and A.R-E. performed split GFP studies, T.M.S, M.M., M.P., J.O.E., B.S., and W.B.F. analyzed the data; T.M.S., M.M., M.P., J.O.E., G.V.D, R.S. and W.B.F. wrote the manuscript.

## Supplementary Materials

**Figure 1-figure supplement 1.**
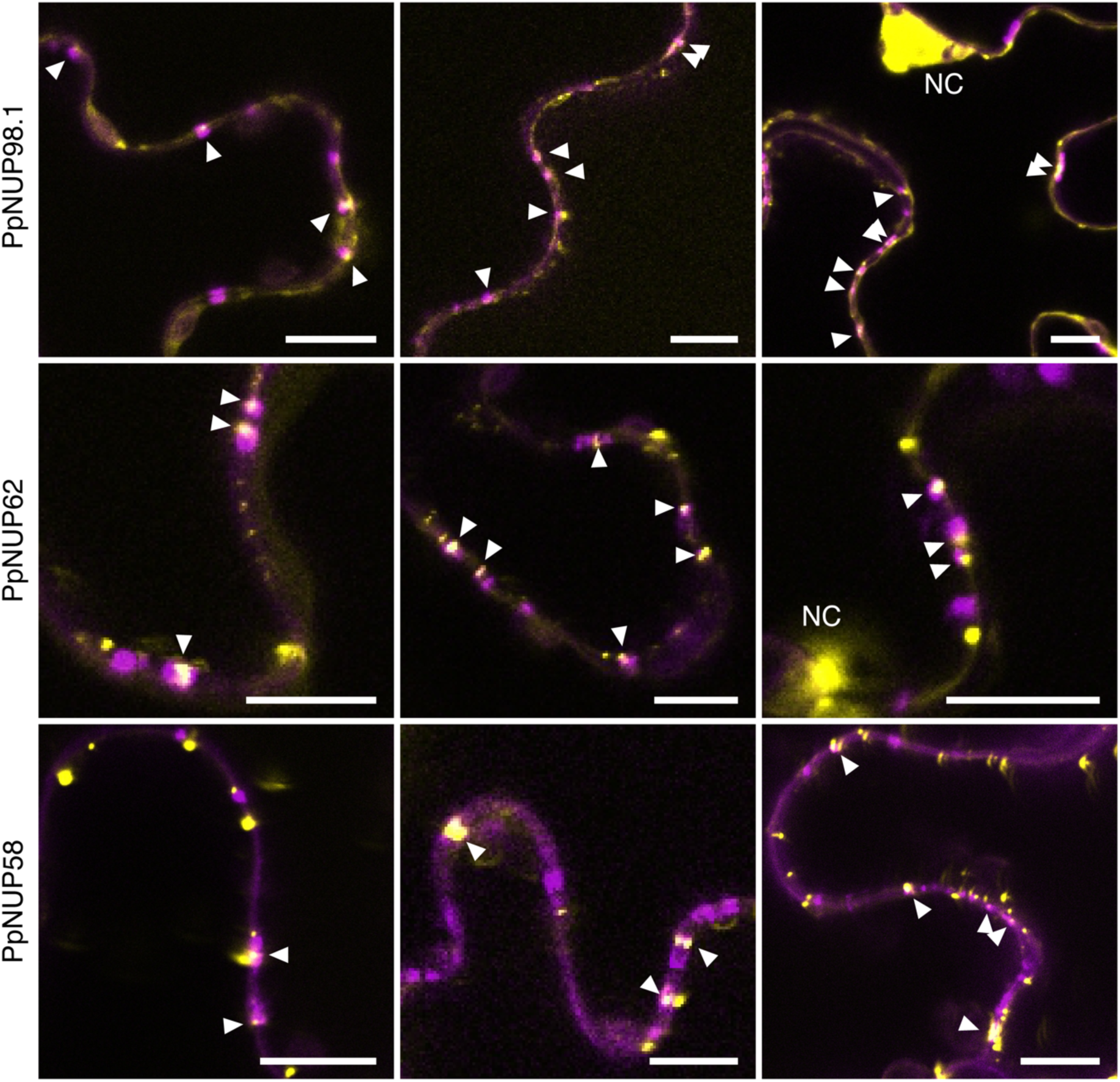
Overlay of *Physcomitrium patens* FG-NUPs with the PD marker aniline blue. Localization of FG-NUPs from *P. patens* transiently expressed in *N. benthamiana* epidermal cells visualized by a single confocal optical section. FG-NUPs were fused at their C-terminus to mVenus (yellow) and expressed transiently under the control of the inducible XVE promotor. Aniline blue (purple) was infiltrated to stain callose, which accumulates in pit fields. Arrowheads indicate overlay between NUPs and aniline blue. For each NUP, a similar localization pattern appeared in at least three independent experiments. NC = nucleus. Scale bar: 10 µm.

**Figure 2-figure supplement 1.**
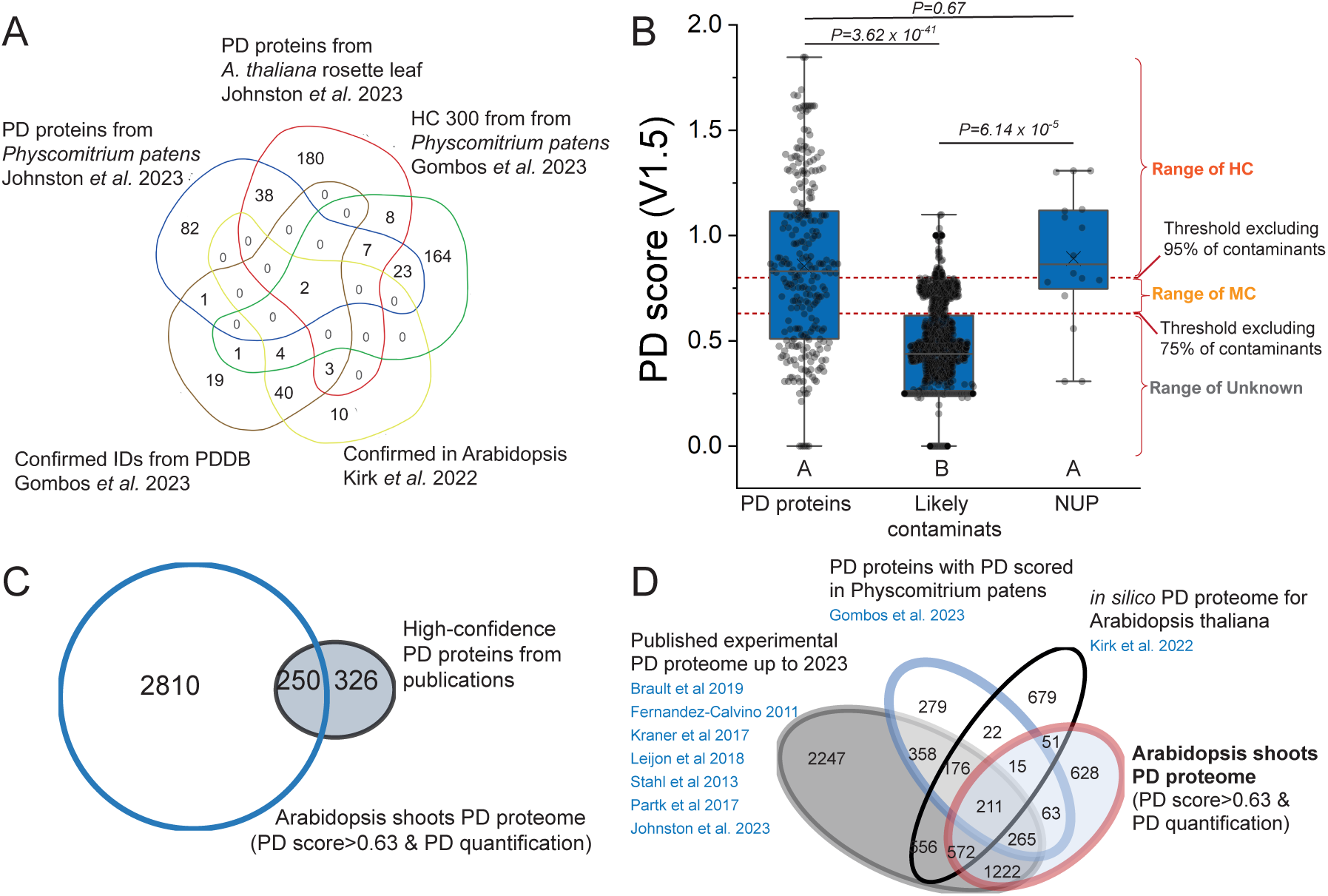
Arabidopsis shoot proteome and the comparison to other PD proteomes. (**A)**, Venn diagram defining high-confidence PD proteins from previous publications, e.g., confirmed Arabidopsis PD proteins, and high confidence PD proteins identified in *Physcomitrium*, and in both *Physcomitrium* and Arabidopsis leaves (Gombos et al., 2023; Johnston et al., 2023; Kirk et al., 2022). These proteins were then used as high-confidence data set to establish the PD probability threshold and were labelled as ’PD proteins’ in file S3. 43% of these proteins were quantified in our Arabidopsis shoot PD proteome. (**B)**, PD Scores of nuclear pore components. PD Scores (v 1.5) are shown as box plots for high-confidence PD proteins (PD proteins), typical co-purifying proteins from plastid and mitochondria identified in Arabidopsis PD preparations (likely contaminants) and proteins of the nuclear pore complex (NUP). Dotted lines indicate two thresholds based on the frequency distribution for contaminant proteins and PD proteins. The median is shown as a gray line, the mean value is denoted by a + symbol, and the whiskers cover the 95^th^percentile. PD Scores above 0.83 (the upper dotted line) indicate high-confidence (HC) PD proteins; scores between 0.63 and 0.83 (between the two dotted lines) signify medium-confidence (MC) PD proteins; and scores below 0.63 are considered low confidence (LC) as their PD Scores overlapped with 75% of the contaminants. These confidence clusters were annotated in file S3. The final Arabidopsis shoot PD proteome consists of proteins with PD scores above 0.63, which are also quantifiable in the PD fraction. Capital letters represent group statistical comparisons determined by one-way ANOVA. The p-values of pairwise comparison using Student’s t-test, are indicated on the top of the boxes. (**C)**, Overlap between the Arabidopsis shoot PD proteome and other PD proteomes. Proteins with a PD score >0.63 and quantifiable in PD fraction (Arabidopsis shoot PD proteome) were compared with nine representative PD proteome studies. In the current Arabidopsis shoot PD proteome, 80% of the proteins were previously identified in published PD proteomes. Combining PD enrichment and data analysis thresholding allowed us to determine the confidence levels of PD proteins and obtain a “cleaner” PD proteome. In total, we identified 5145 proteins from PD, CW, and TC fractions, with 3347 proteins having a PD score >0.63. Among them, 3062 proteins are quantifiable in the PD fraction, constituting our Arabidopsis shoot PD proteome. Among these, 2268 proteins are categorized as high-confidence PD proteins, while 792 proteins fall into the medium-confidence category (file S3). Additionally, we discovered 20 nuclear pore proteins (NUPs), with 15 of them surpassing the threshold and being included in the final PD proteome. Among these, 11 are classified as high-confidence PD proteins, while 4 are medium-confidence PD proteins (file S3, Figure 1-figure supplement 2B). (**D)** Comparison of Arabidopsis shoot PD proteomes with various published PD proteomes. 80% of the proteins had been identified in previous studies.

**Figure 2-figure supplement 2.**
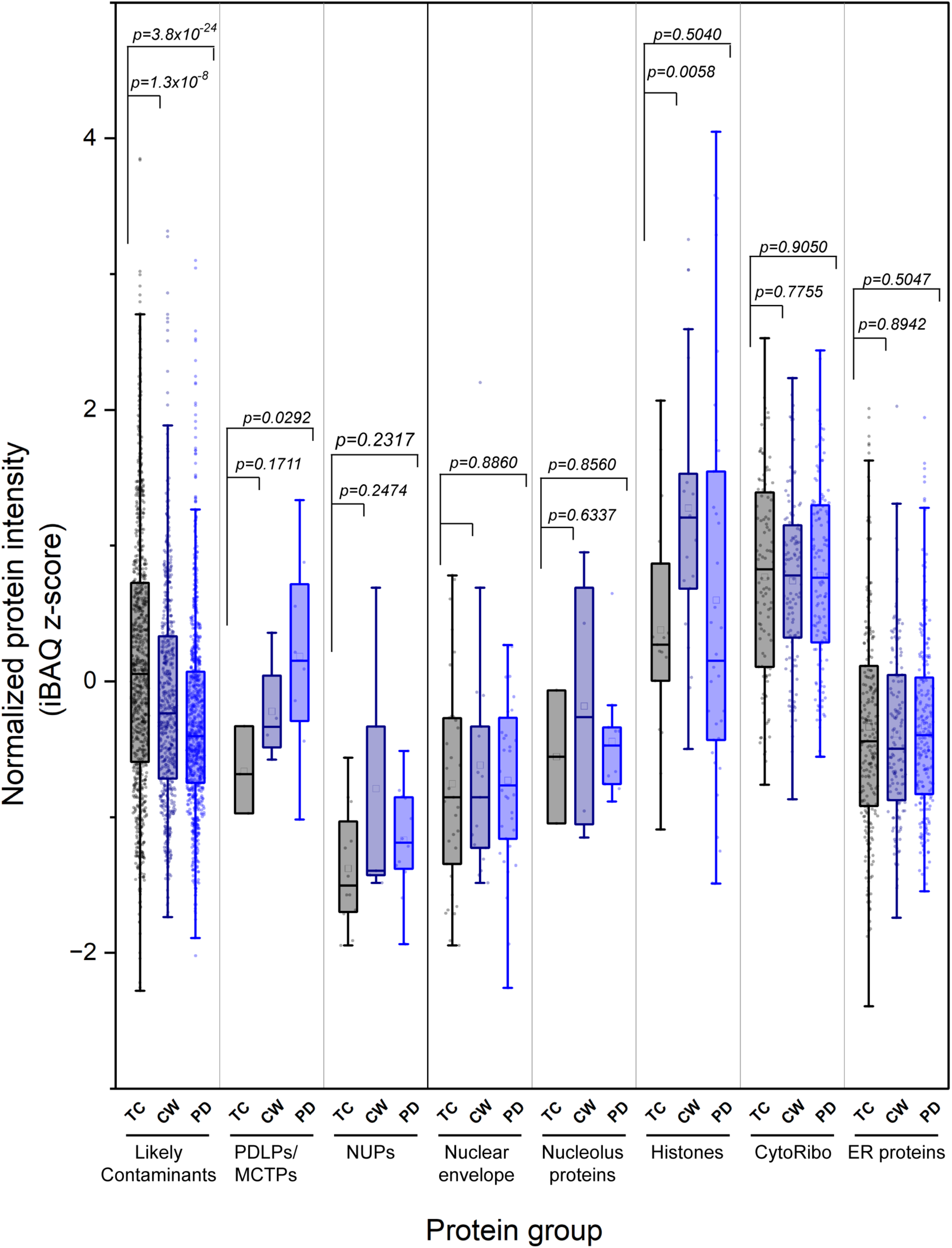
Normalized protein intensities of different protein groups in total extract (TC), cell wall fraction (CW) and PD enrichment (PD). The group of likely contaminants contains plastidic and mitochondrial proteins. *Bona fide* PD proteins are PDLPs and MCTPs. Protein categorization was based on Mapman (Usadel et al., 2005). A Student’s t-test was used for statistical analysis.

**Figure 3-figure supplement 1.**
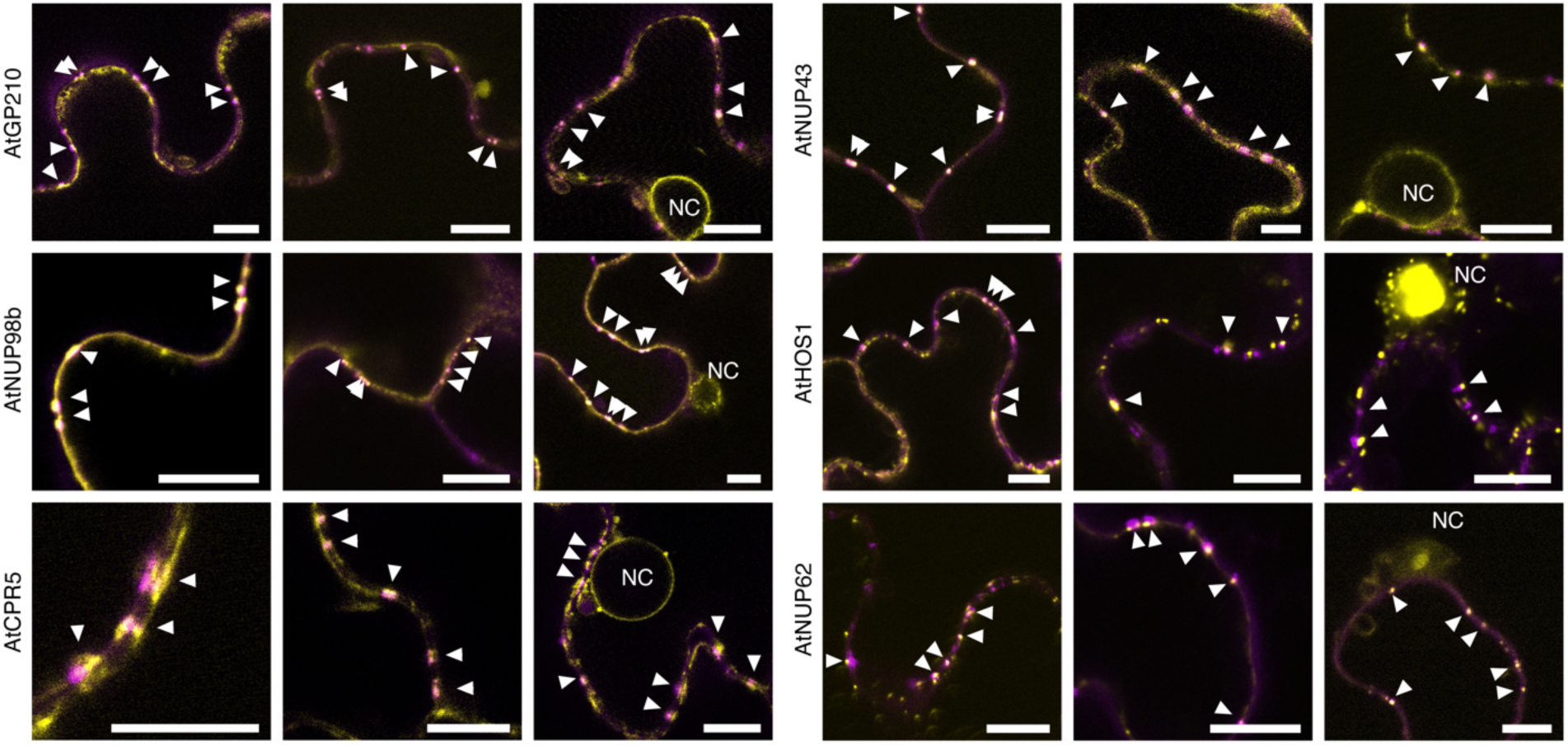
Arabidopsis NUP-FP localization overlays with a PD marker. Localization of Arabidopsis NUPs transiently expressed in *N. benthamiana* epidermal cells visualized by a single confocal optical section. NUPs were fused at their C-terminus to mVenus or mCitrine (yellow) and expressed transiently under the control of the inducible XVE promotor. Aniline blue (purple) was infiltrated to stain callose that accumulates in pit fields. Arrowheads indicate overlay between NUPs and aniline blue. NUPs shown here were found to have a significant increased PD index values compared to negative controls in Figure 2. For each NUP, the localization pattern was reproducible in at least in three independent experiments. NC, nuclei. Note that for comparison, we here show the same images for NUP98B and NUP62 from Figure 2A again. Scale bar: 10 µm.

**Figure 5- figure supplement 1.**
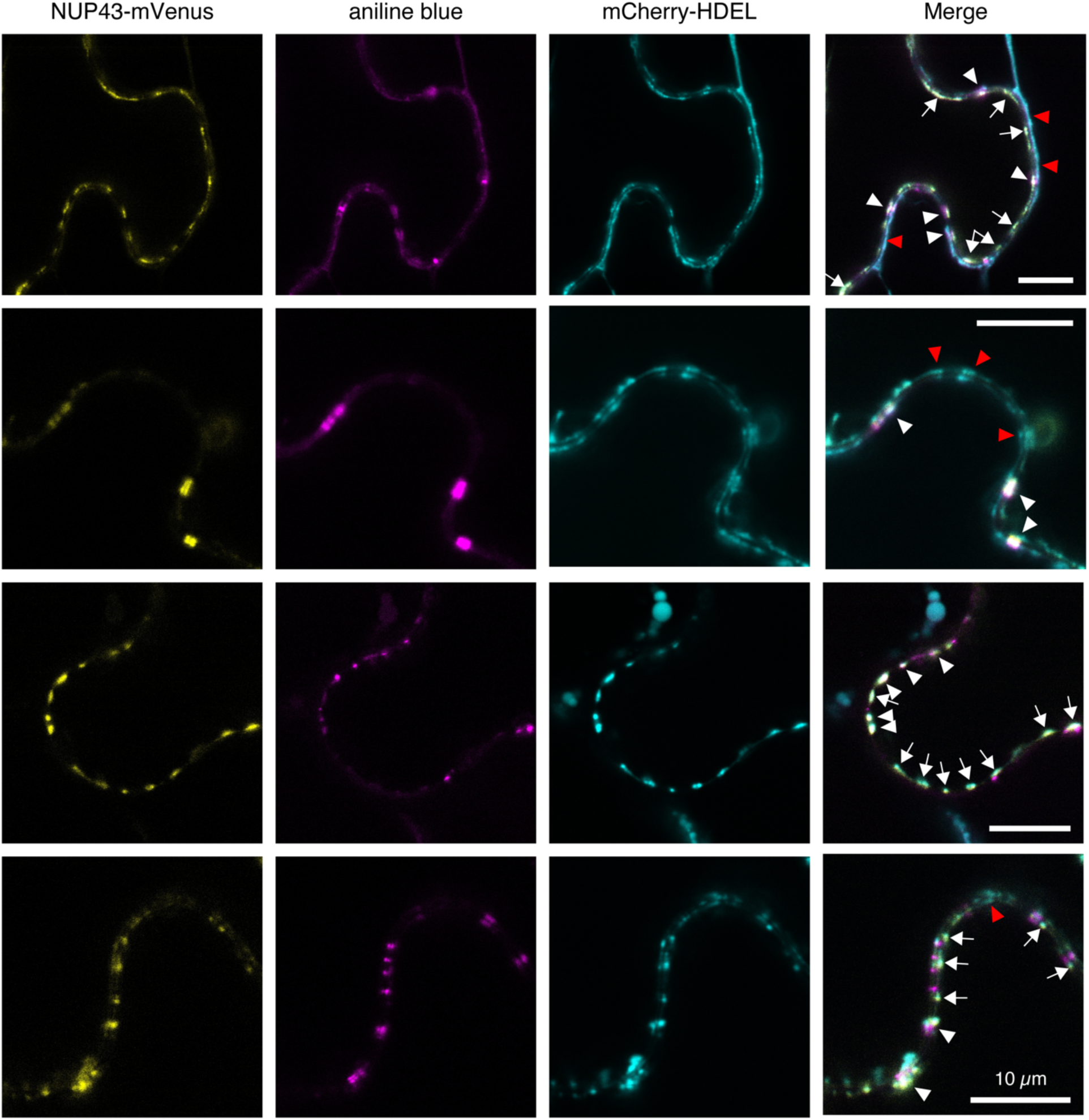
Arabidopsis NUP43 partially co-localizes with ER marker. Transient co-expression of NUP43-mVenus and mCherry-HDEL in *N. benthamiana* leaf epidermal cells, visualized by a single confocal optical section. Aniline blue was infiltrated to detect callose in pit fields as PD marker. White arrowheads indicate overlay of NUP43-mVenus, mCherry-HDEL. And aniline blue. Red arrows indicate mCherry-HDEL specific localization., without overly with other markers. White arrows (*middle right* panel) indicate overlay of NUP43-mVenus and mCherry-HDEL but absent aniline blue staining. Scale bar indicates 10 µm.

**Figure 6-figure supplement 1.**
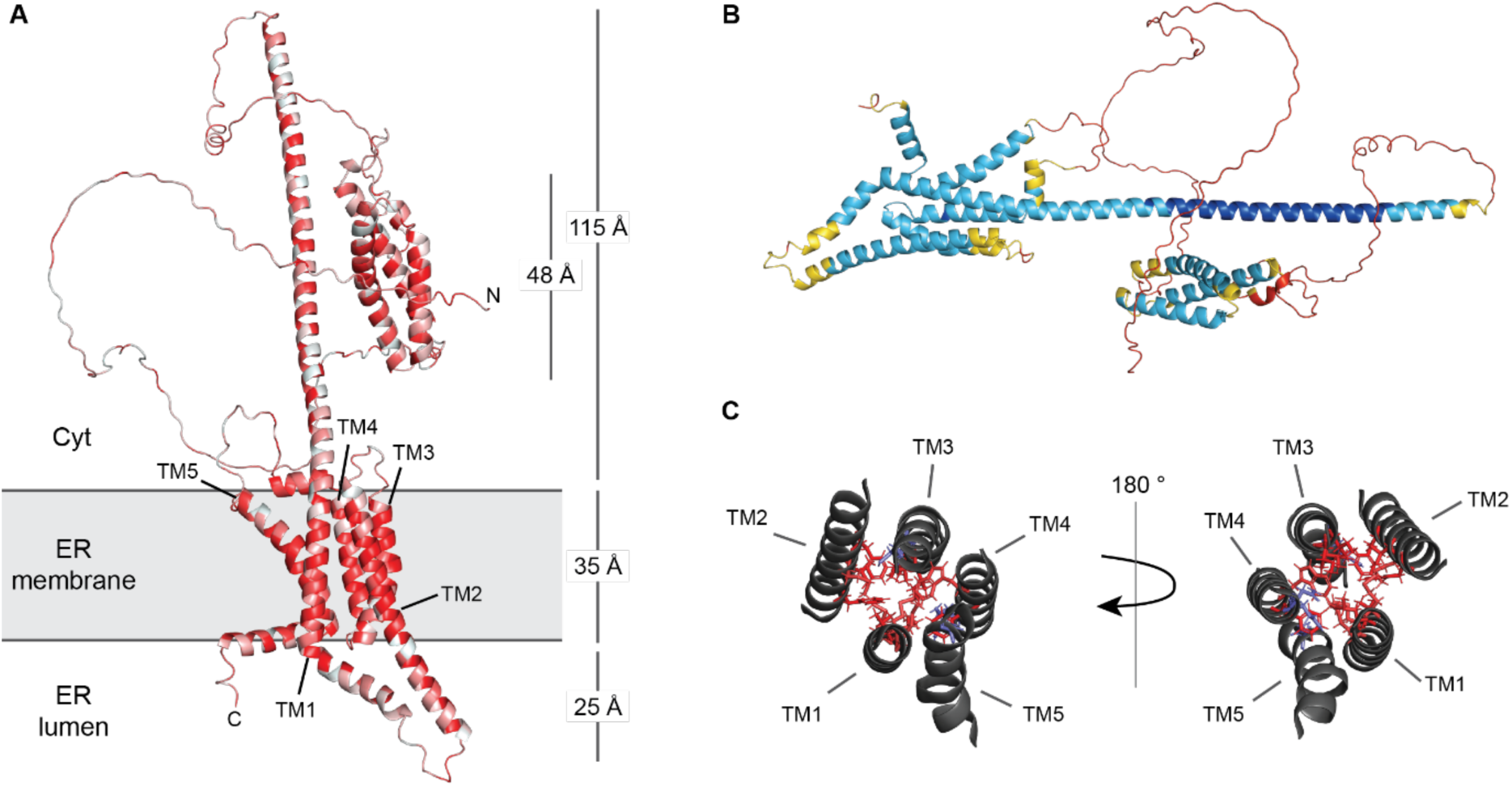
Predicted structure of monomeric CPR5 and model for integration into ER-derived membranes. Topological studies using the split-GFP system (Figure 4A,B, Figure 4-figure supplement 3) are consistent with a localization of the CPR5 C-terminus in the ER lumen, while its N-terminus extends into the cytoplasm. (**A)**, CPR5 is composed of a N-terminal, disordered region (residues 1-106), a soluble domain formed by 3 α-helices (107-192), linked by a second disordered region (193- 265) to an approx. 115 Å long, central α-helix (266-342). Hydrophobic residues (red) of the central helix (TM1) and 4 shorter helices (TM2-TM5) form a transmembrane domain spanning approx. 35 Å, consistent with the expected thickness for ER-derived membranes. C-terminus and helical extensions of TM1 and TM2 reach into the ER lumen. (**B)**, Structure colored by per-residue prediction confidence (pLDDT; dark blue: 90-100, light blue: 50-70, yellow: 50-70, red: < 50). Low confidence values (red) may indicate regions that are either intrinsically disordered or assume a structure when interacting with other proteins and therefore could partake in formation of complexes with NUPs. (**C)**, View of transmembrane helical bundle from ER lumen (left) and cytoplasm (right). Amino acid side chains spanning between transmembrane helices are colored by hydrophobicity (blue: hydrophilic, red: hydrophobic). The CPR5 transmembrane domain is not predicted to form a solvent-accessible pore.

**Figure 6-figure supplement 2:**
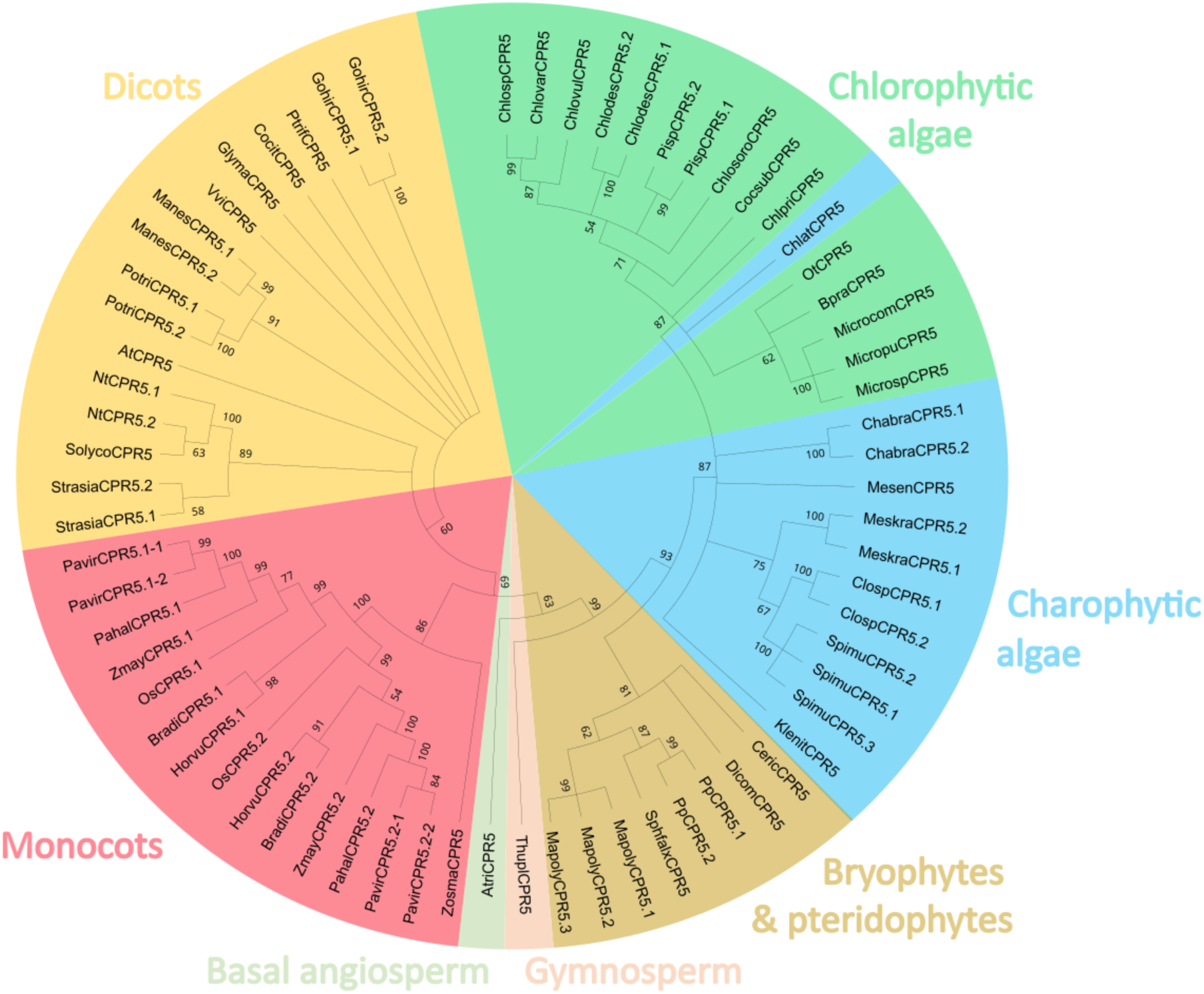
Phylogenetic analysis of CPR5 sequences. The accession numbers and the abbreviations corresponding to the mature protein sequences used for this analysis are given in table S7. The analysis was performed as described in the “Materials and Methods” section.

**Figure 6-figure supplement 3.**
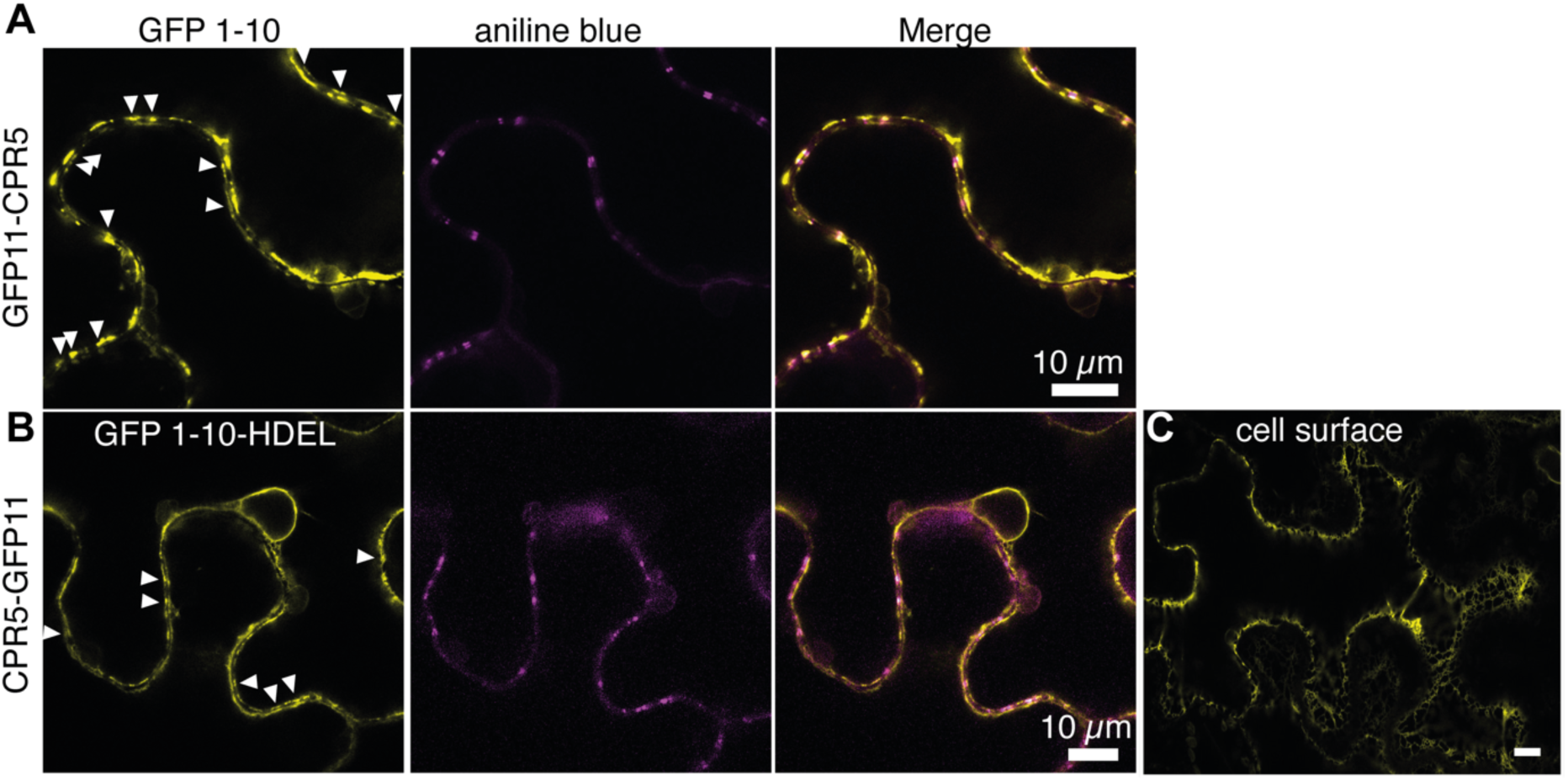
The C-terminus of Arabidopsis CPR5 faces ER lumen. **(A)**, Transient co-expression of GFP11-CPR5 and cytosolic GFP1-10 in *N. benthamiana* leaves, same cell as shown in Figure 4*. Left*, GFP channel. Reconstitution can be observed also at PD (arrowheads). *Middle*, aniline blue*. Right,* merge. (**B)**, *N. benthamiana* epidermal cells transiently expressing CPR5-GFP11 and ER luminal GFP1-10-HDEL, same cell as shown in Figure 4*. Left*, GFP channel. Reconstitution can be observed also at PD (arrowheads); *middle*, aniline blue*; right,* merge. (**C)**, Surface view of the same cell as in B, showing reconstitution in the ER. Confocal images in Figure 6-figure supplement 3 are single optical sections.

**Figure 6-figure supplement 4.**
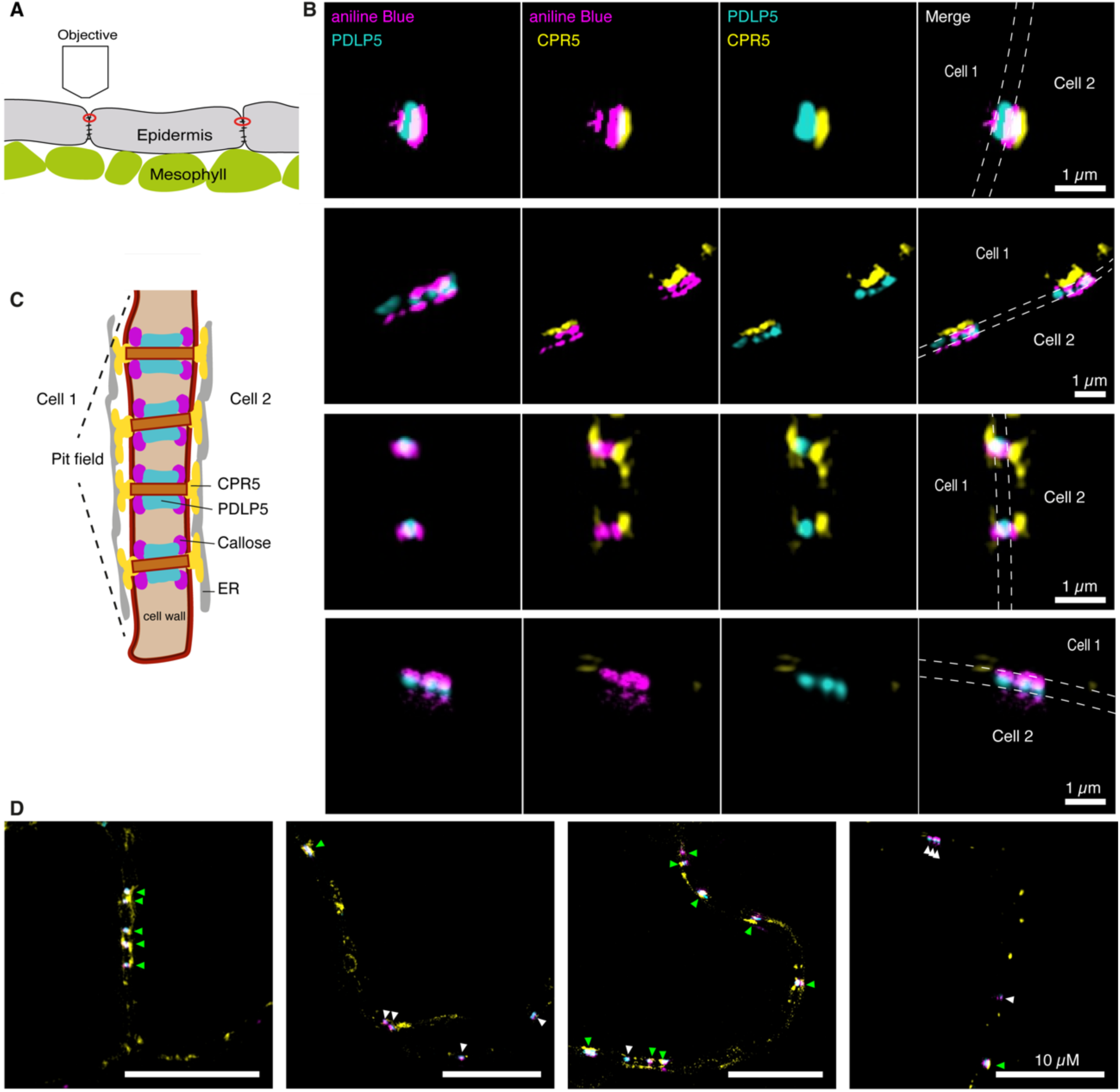
**Arabidopsis CPR5-mCitrine localizes at the orifices of PD**. (**A),** Cartoon showing analyzed PD: only PD most proximal to the cover glass and objective were analyzed to reduce effects caused by scattering of light due to cell wall. (**B),** single optical sections recorded using Structured Illumination Microscopy (SIM) in *N. benthamiana* leaves co-infiltrated with XVE:PDLP5-mScarlet3 and XVE:CPR5-mCitrine. Prior to the imaging, leaves were infiltrated with aniline blue. PDLP5-mScarlet3 in turquoise, CPR5-mCitrine in yellow, and aniline blue in magenta. Comparable results were obtained in four independent experiments. Dashed lines indicate approximation of cell wall boundaries. (**C),** Cartoon of pit field and localization of PDLP5 and CPR5. Fluorescence reporting on CPR5 in (B) extends laterally 200-1000 nm in the epidermal cell wall. Previously, the lateral extension of individual PD pores in the cell wall was estimated to 100 nm for basal trichome cell walls and 187 nm for epidermal cell walls (Faulkner et al., 2008; Fitzgibbon et al., 2010). The resolution achieved with SIM was not high enough to resolve single PD or determine whether pit fields consist of several simple PD (as shown in C) or a few complex (H-shaped/twinned) PD. (**D**), single optical sections recorded using SIM in *N. benthamiana* leaves co-infiltrated with XVE:PDLP5-mScarlet3 and XVE:CPR5-mCitrine. Green arrowheads indicate overlay between CPR5-mCitrine, PDLP5-mScarlet3 and aniline blue; white arrowheads indicate overlay between PDLP5-mScarlet3 and aniline blue, without CPR5-mCitrine. PDLP5-mScarlet3 in turquoise, CPR5-mCitrine in yellow, and aniline blue in magenta.

**Figure 7-figure supplement 1.**
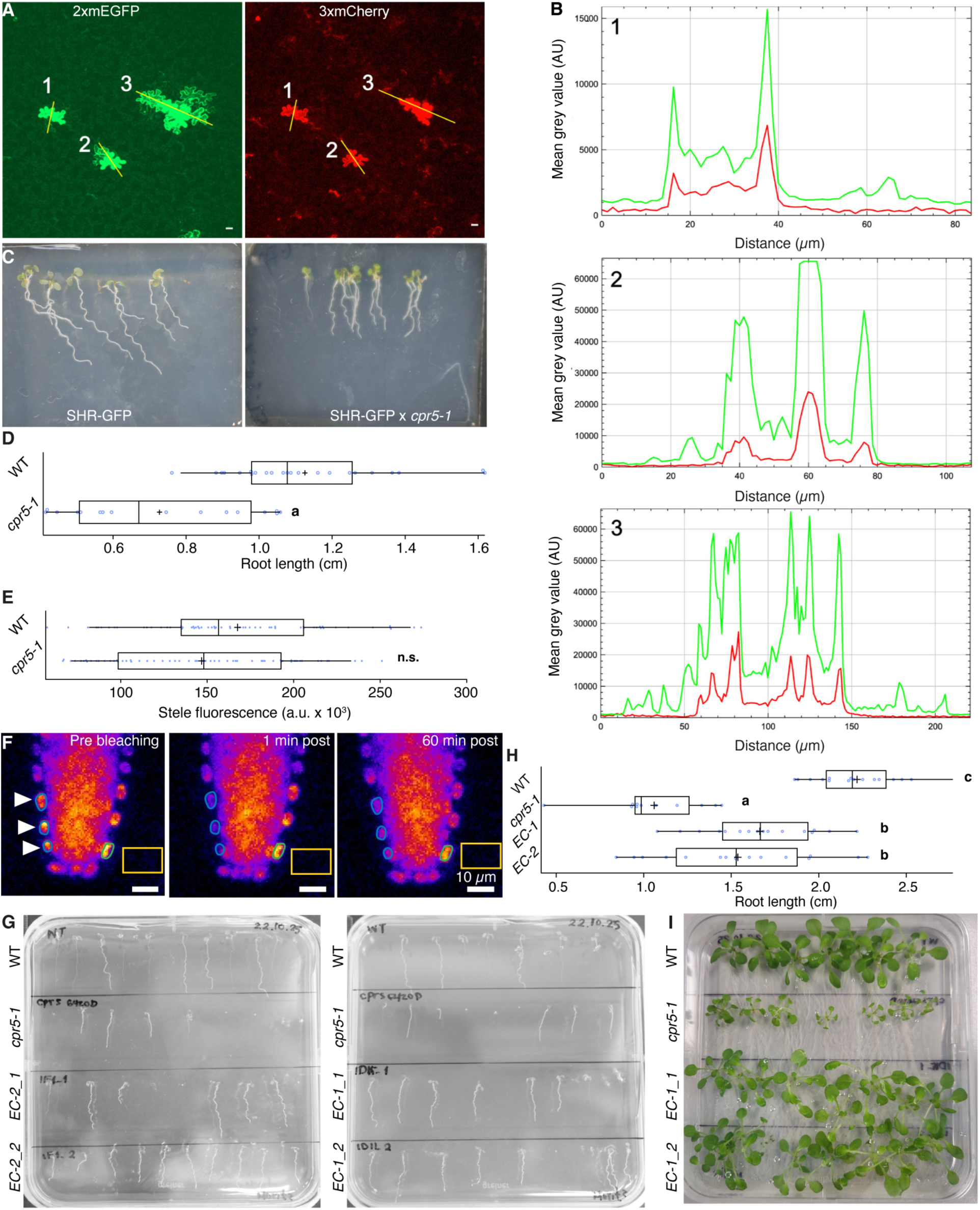
Intercellular transport assays in *cpr5-1* mutants. **(A)**, Example of maximum intensity projection of 32 optical cross-sections taken at 2.5 µm intervals with three different clusters (1, 2, and 3) of fluorescent cells after bombardment in a WT Arabidopsis leaf. Each cluster reports a successful bombardment event. Left, GFP channel, right, mCherry channel. 24 h prior to the experiment, leaves were co-bombarded with gold particles coated with p35S:mEGFP-mEGFP (2xmEGFP). DNA plasmids and gold particles coated with p35S:mCherry-mCherry-mCherry (3xmCherry). (**B),** Profile plots across the different clusters, 1, 2, and 3 as shown in A. Yellow lines in A indicate the position for the profile. The bombarded cell marked with 3xmCherry overlays with the brightest cell in the GFP channel. (**C),** Six-day old Arabidopsis seedlings with *shr*/pSHR:SHR-GFP in WT Col-0 background *(left)* and *shr*/pSHR:SHR-GFP in the *cpr5-1* background (*right*) grown on ½ MS agar. **(D),** Quantification of the root length of 6-day old WT and *cpr5-1* mutant background seedlings. n_(WT)_ = 23 roots, four independent experiments, n_(*cpr5-1)*_ *=* 16 roots, three independent experiments. **a** Significantly decreased root length in *cpr5-1* mutant, based on Student’s *t*-test with *p* = 0.00002. **(E),** Quantification of SHR-GFP stele baseline fluorescence in *cpr5-1* mutant and WT background. n_(WT)_ = 62 stele ROIs, n_(*cpr5-1*)_ = 62 stele ROIs, with 3-4 ROIs analyzed per root. **n.s.,** no significant difference in fluorescence intensity based on Mann-Whitney U test with *p* = 0.074. **(F)**, SHR–GFP fluorescence recovery after photobleaching (FRAP) in homozygous *shr*/pSHR:SHR-GFP root endodermal cells (arrow heads). Note that for comparison, we here show the same sum projections as in Figure 5F (*top*). Blue circles show the position of ROIs for FRAP. Green circle shows the position of ROI for control nucleus used for normalization (see Methods). Yellow square shows the position for mean background fluorescence used for correction (see Methods). **(G)**, Seven-day old Arabidopsis WT Col-0 and *cpr5-1* mutant seedlings (*left* and *right*), and *cpr5-1*/pCPR5:CPR5-mCitrine independent expression complementation (EC) lines (*EC-1 and EC-2*). For each complementation line, two sublines are shown (*left*: *EC-1_1* and *EC-1_2*; *right: EC-2_1, EC-2_2*), grown on ½ MS agar. **(H),** Quantification of the root length of 7-day old WT and *cpr5-1* mutant background, as well as pooled *EC-1* lines and *EC-2* lines. Quantification is based on two independent experiments with n_(WT)_ = 16 roots, n_(*cpr5-1)*_ *=* 11 roots, n_(*IF_1*)_ = 16 roots, n_(*EC-1*)_ = 16 roots. Different letters indicate significant difference tested by one-way ANOVA (p = 5 x 10^-11^) and Tukey *post hoc* analysis. p(_WT vs. *cpr5-1*_) = 2 x10^-11^, p(*_EC-2_* _vs. *cpr5-1*_) = 0.003, p(*_EC-1_* _vs *cpr5-1*_) = 0.0001, p(_WT vs *EC-2*_) = 0.000002, p(_WT vs *EC-1*_) = 0.0001. **(I),** same plate as in (G) *right* but 3 weeks later, with seedlings grown 4-weeks in total.

## Supplementary Tables

**table S1.**
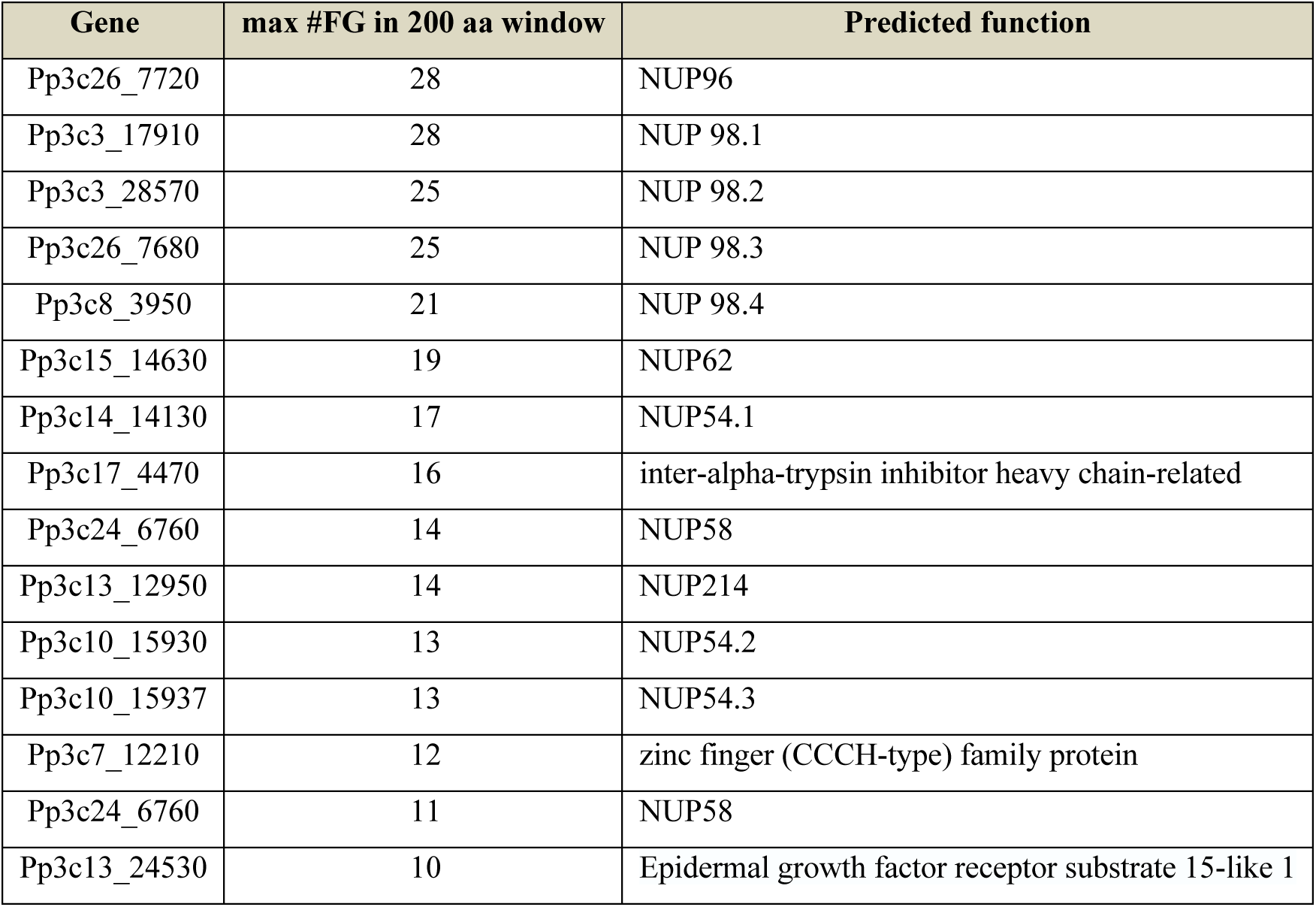
Genome-wide search for potential phase-separating proteins with FG repeats. . The 15 genes with the highest number of FG motifs within a 200-amino acid window in the *Physcomitrium* genome. For the list summarizing the genome-wide search for FG containing genes, see file S1. For Arabidopsis NUPs see file S2.

| Gene | max #FG in 200 aa window | Predicted function |
| --- | --- | --- |
| Pp3c26_7720 | 28 | NUP96 |
| Pp3c3_17910 | 28 | NUP 98.1 |
| Pp3c3_28570 | 25 | NUP 98.2 |
| Pp3c26_7680 | 25 | NUP 98.3 |
| Pp3c8_3950 | 21 | NUP 98.4 |
| Pp3c15_14630 | 19 | NUP62 |
| Pp3c14_14130 | 17 | NUP54.1 |
| Pp3c17_4470 | 16 | inter-alpha-trypsin inhibitor heavy chain-related |
| Pp3c24_6760 | 14 | NUP58 |
| Pp3c13_12950 | 14 | NUP214 |
| Pp3c10_15930 | 13 | NUP54.2 |
| Pp3c10_15937 | 13 | NUP54.3 |
| Pp3c7_12210 | 12 | zinc finger (CCCH-type) family protein |
| Pp3c24_6760 | 11 | NUP58 |
| Pp3c13_24530 | 10 | Epidermal growth factor receptor substrate 15-like 1 |

**table S2.** List of known Arabidopsis NUPs and summary of localization results. *p*-values calculated from Bonferroni-corrected pairwise comparison between the tested NUP with propidium iodide marked by *; *p*-values calculated from Bonferroni-corrected pairwise comparison between the tested NUP and free mScarlet3 marked by #. Index is short for PD index (see also Figure 2 and 3). NA: not assessed. FG-NUPs in red.

| NUP | AGI | Localization at NPC | Tested for localization |
| --- | --- | --- | --- |
| NDC1 | AT1G73240 | membrane anchor | NA |
| Gp210 | AT5G40480 | membrane anchor | index = 1.77, * = $2 \times 10^{-9}$; # = $3 \times 10^{-6}$ |
| CPR5 | AT5G64930 | membrane anchor | index = 1.48, * = $7 \times 10^{-8}$; # = **0.0001** |
| PNET1 | AT1G07970 | membrane anchor | NA |
| NUP155 | AT1G14850 | inner ring | index = 1.34, * = **0.002**; # = 1.0 |
| NUP93a | AT2G41620 | inner ring | index = 1.38, * = **0.001**; # = 1.0 |
| NUP35/MPPN | AT3G16310 | inner ring | index = 1.36, * = **0.0002**; # = 0.549 |
| NUP93b | AT3G57350 | inner ring | index = 1.19, * = 0.43; # = 1.0 |
| NUP188 | AT4G38760 | inner ring | NA |
| NUP205 | AT5G51200 | inner ring | NA |
| NUP160 | AT1G33410 | outer ring | NA |
| Seh1 | AT1G64350 | outer ring | NA |
| NUP96 | AT1G80680 | outer ring | NA |
| NUP133 | AT2G05120 | outer ring | NA |
| Sec13 | AT2G30050 | outer ring | NA |
| Elys/HOS1 | AT2G39810 | outer ring | index = 1.52, * = $3 \times 10^{-7}$; # = **0.001** |
| NUP107 | AT3G14120 | outer ring | NA |
| NUP43 | AT4G30840 | outer ring | index = 1.72, * = $3 \times 10^{-6}$; # = **0.006** |
| NUP75/85/SBB1 | AT4G32910 | outer ring | NA |
| NUP98a | AT1G10390 | core FG-NUP | index = 1.38, * = **0.006**; # = 1.0 |
| NUP54 | AT1G24310 | core FG-NUP | NA |
| NUP98b | AT1G59660 | core FG-NUP | index = 1.55, * = **0.0001**; # = 0.276 |
| NUP62 | AT2G45000 | core FG-NUP | index = 1.34, * = **0.0001**; # = 0.357 |
| NUP58 | AT4G37130 | core FG-NUP | index = 1.37, * = **0.001**; # = 1.0 |
| CG1 | AT1G75340 | cytoplasmic FG-NUP | NA |
| GLE1 | AT1G13120 | cytoplasmic NUP | NA |
| NUP214 | AT1G55540 | cytoplasmic FG-NUP | index = 1.29, * = **0.041**; # = 1.0 |
| Rae1 | AT1G80670 | cytoplasmic NUP | index = 1.22, * = 0.301; # = 1.0 |
| Aladin | AT3G56900 | cytoplasmic NUP | NA |
| MOS7/NUP88 | AT5G05680 | cytoplasmic NUP | NA |
| NUP50a | AT1G52380 | basket FG-NUP | NA |
| NUA/TPR/ATTPR | AT1G79280 | basket | NA |
| NUP1/NUP136 | AT3G10650 | basket FG-NUP | NA |
| NUP50b | AT3G15970 | basket FG-NUP | index = 1.06, * = 1.0; # = 1.0 |
| NUP82 | AT5G20200 | basket | index = 1.24, * = 0.1; # = 1.0 |

**table S4.** NPC components identified in published cell wall/PD proteomes.

| **NUP** | **Reference** |
| --- | --- |
| CPR5 | (Kraner et al., 2017) |
| NUP93a | (Kirk et al., 2022) |
| NUP155 | (Bayer et al., 2006; Fernandez-Calvino et al., 2011; Kraner et al., 2017) |
| SEC13 | (Bayer et al., 2006; Gombos et al., 2023; Kraner et al., 2017) |
| NUP205 | (Kirk et al., 2022) |
| NUA | (Gombos et al., 2023) |
| NUP50a * | (Gombos et al., 2023; Kraner et al., 2017; Li et al., 2024) |
| GP210 | (Kraner et al., 2017) |
| NUP35 | (Kraner et al., 2017) |
| HOS1 | (Kraner et al., 2017) |
| NUP43 | (Kraner et al., 2017) |
| NUP62 | (Kraner et al., 2017) |
| NUP214 | (Kraner et al., 2017) |
| NUP82 | (Kraner et al., 2017) |
| NDC1 * | (Li et al., 2024) |

\* identified by TURBO-ID using PD proteins

**table S5.** Nuclear localization and observation of puncta in the cell periphery, consistent with PD localization by other laboratories.

| NUP | Transient expression | Stable transformation/<br>promoter | Reference |
| --- | --- | --- | --- |
| CPR5 | *p35S:YFP-CPR5* | *p35S:GFP-CPR5/cpr5-1* | (Gu et al., 2016) |
| CPR5* | *p35S:CFP-CPR5* |  | (Peng et al., 2022) |
| NUP98a | *p35S:GFP-Dra2-GFP* | *p35S:NtDra2-GFP* | (Gallemí et al., 2016) |
| NUP155 | *p35S:Nup155-cYFP+p35S:nYFP-CPR5* |  | (Gu et al., 2016) |
| HOS1 | *p35S:HOS1-GFP* |  | (Lee et al., 2012) |
| NUP62 | *p35S:Nup62-nYFP+p35S:FLC-cYFP* |  | (Chen et al., 2023) |

\* split-FP analysis indicated interaction of CPR5 with PRL1 and FIP1

**table S6.**
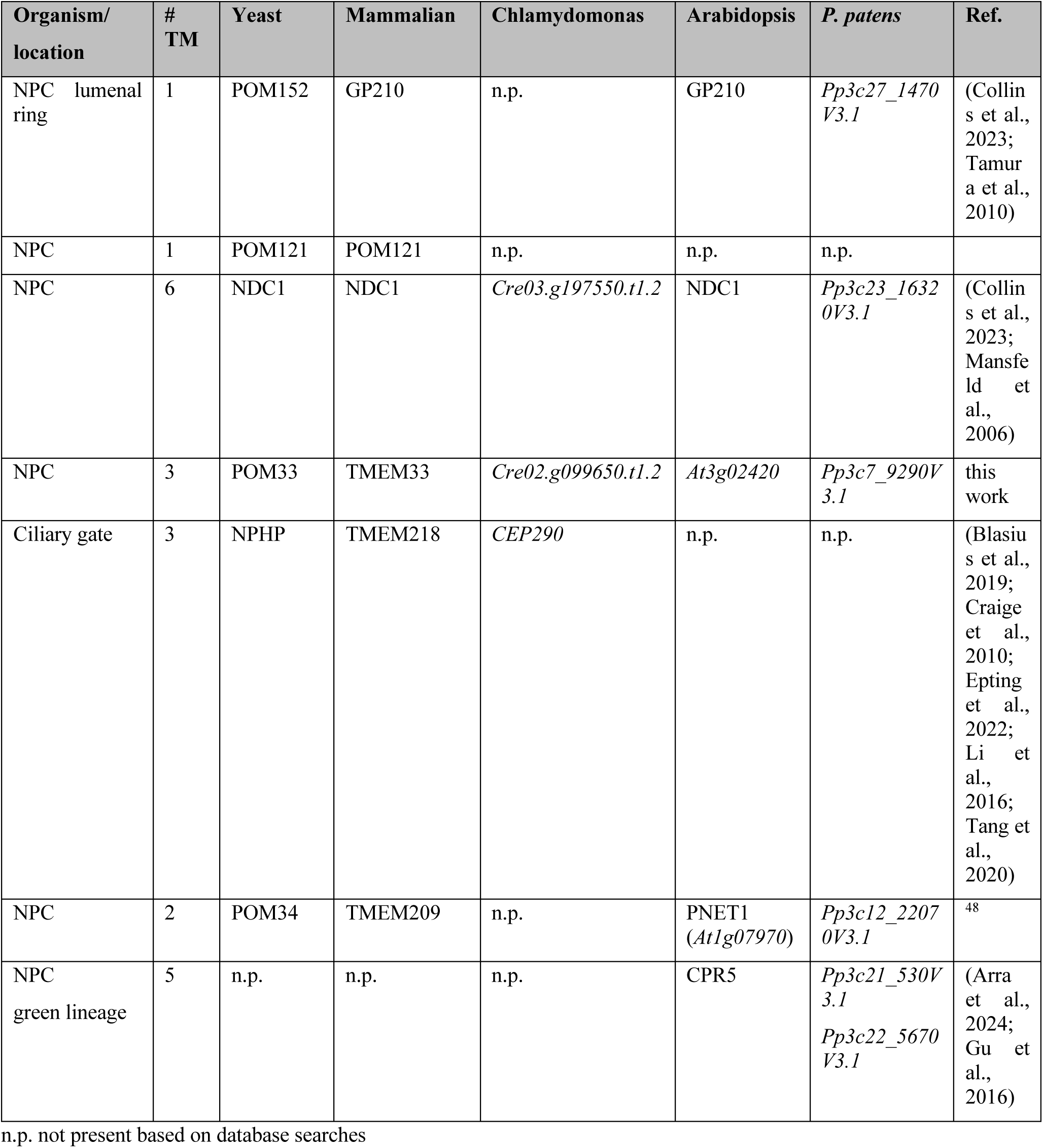
Membrane/microtubule anchor proteins in NPC and ciliary gates across kingdoms.

**table S7.**
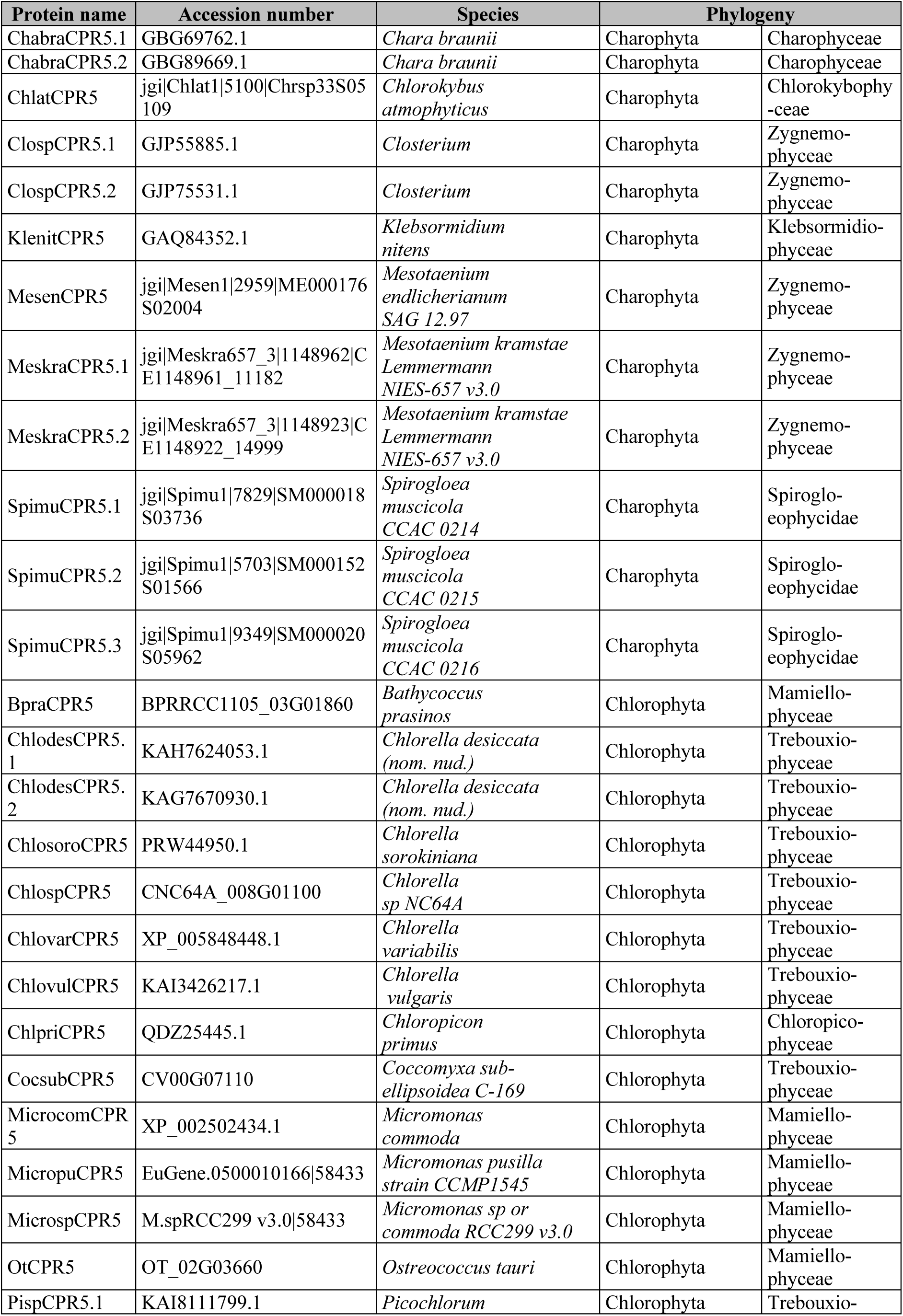
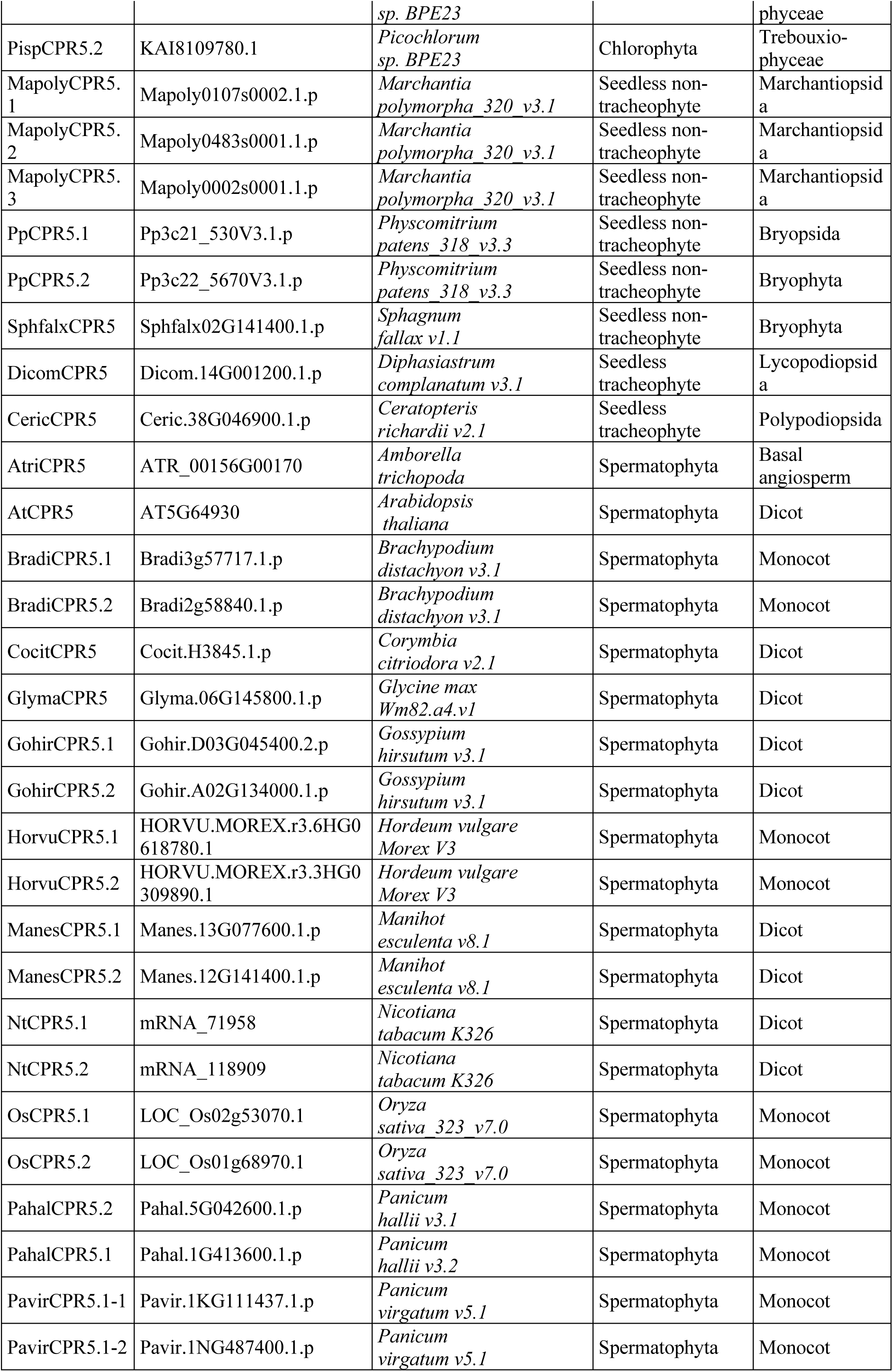
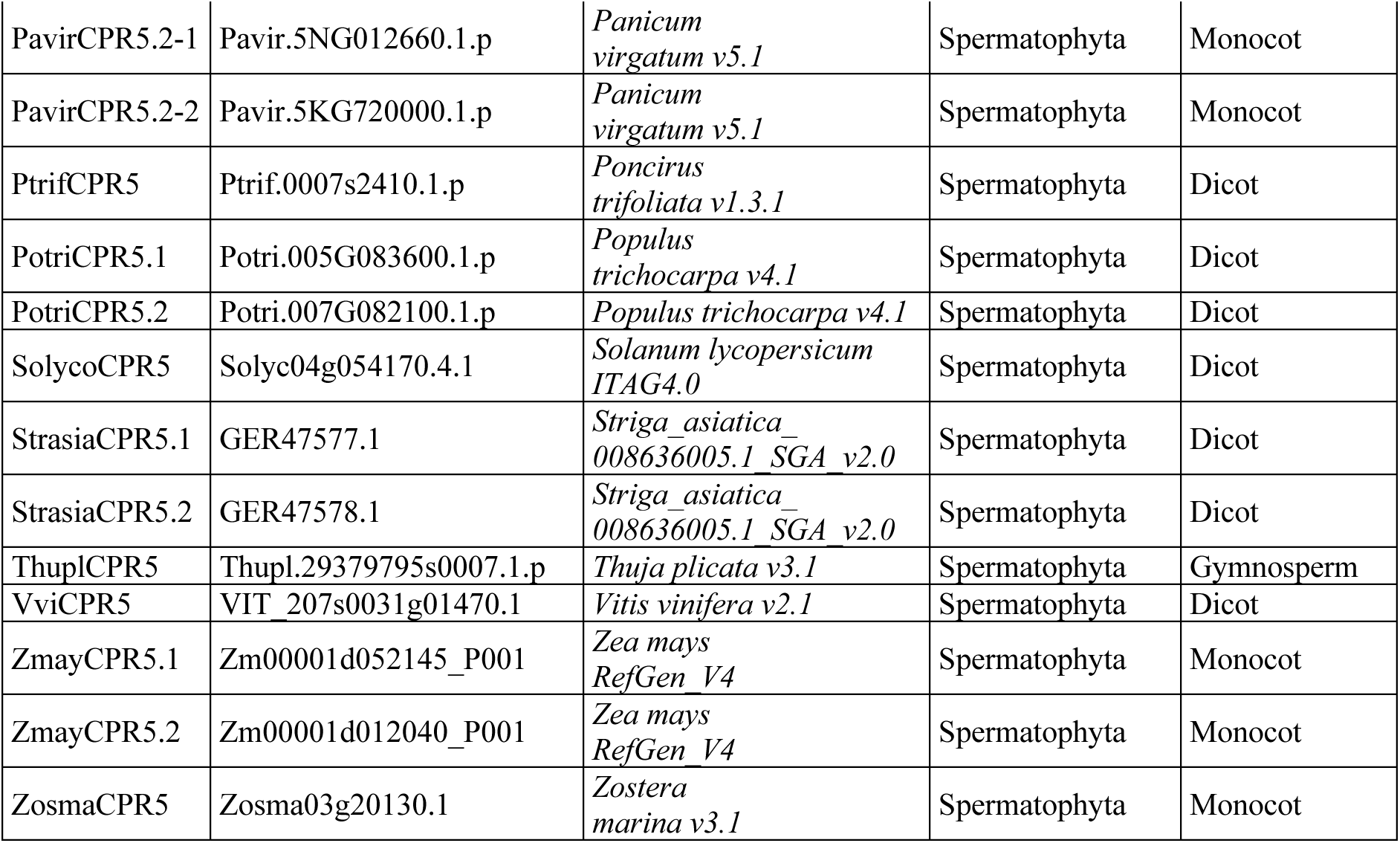
Accession numbers of CPR5 homologs used for the phylogenetic analysis shown in figure S6.

**table S8.**
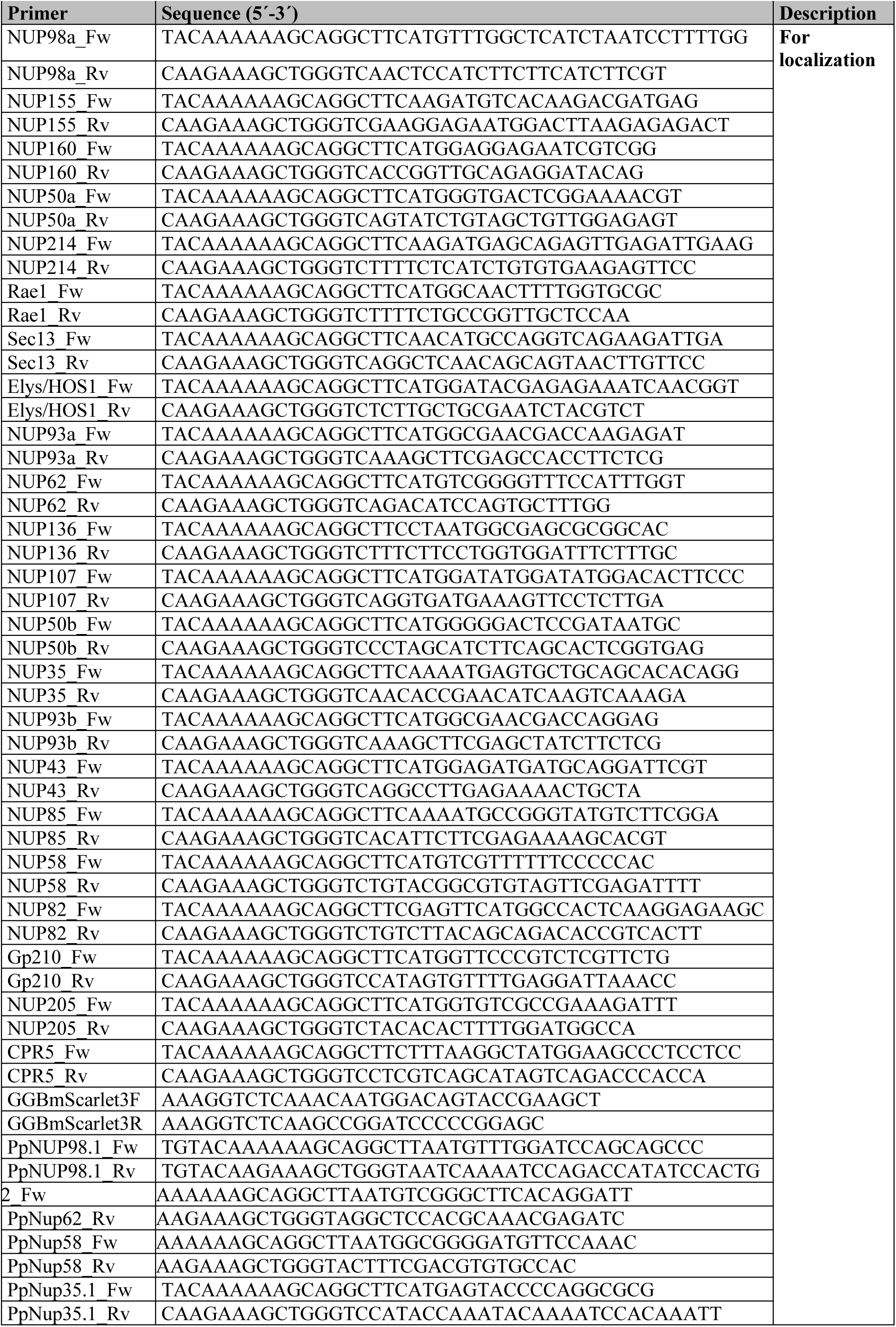
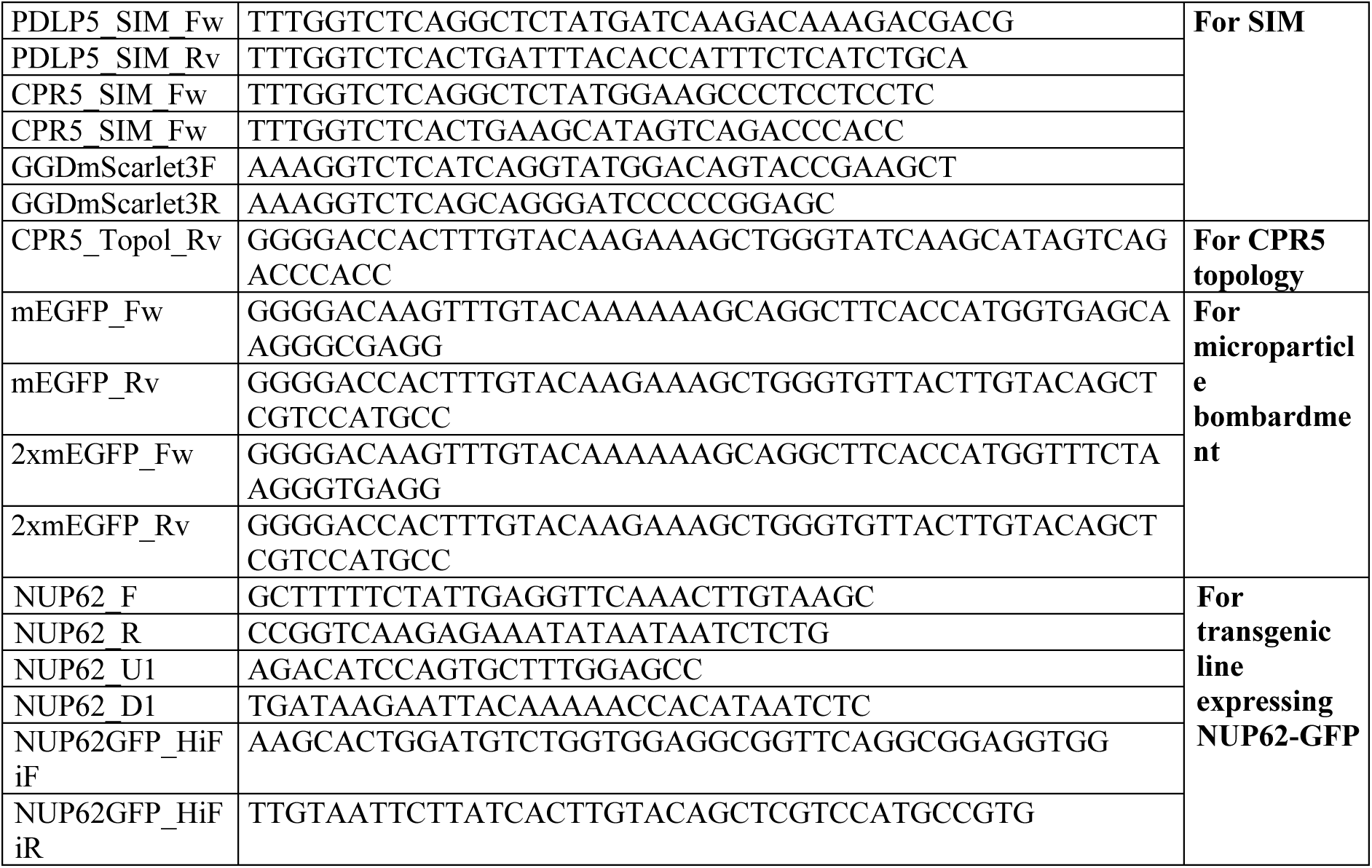
Primers used in this study.

