## Supplementary material for "Identification of nuclear pore proteins at plasmodesmata: potential role in intercellular transport?": Supp Script 2: Supplementary Script 2_Ara.py

```
#!/usr/bin/env python3
import gzip
import io

def read_fasta(file_path):
    sequence_names = []
    sequence_genes = []
    sequences = []

    if file_path.endswith('.gz'):
        with gzip.open(file_path, 'rt', encoding='utf-8') as file:
            process_fasta(file, sequence_names, sequence_genes, sequences)
    else:
        with open(file_path, 'r') as file:
            process_fasta(file, sequence_names, sequence_genes, sequences)

    return sequence_names, sequences, sequence_genes
    
def process_fasta(file, sequence_names, sequence_genes, sequences):
    current_sequence_name = ""
    current_sequence = ""
    for line in file:
        line = line.strip()
        if line.startswith('>'):
            if current_sequence_name:
                sequence_names.append(current_sequence_name.split()[0])
                sequences.append(current_sequence)
                sequence_genes.append(current_sequence_name.split('|')[1])
            current_sequence_name = line[1:]
            current_sequence = ""
        else:
            current_sequence += line
    if current_sequence_name:
        sequence_names.append(current_sequence_name.split()[0])
        sequences.append(current_sequence)
    return sequence_names, sequences, sequence_genes


def count_specific_sequence(sequence, specific_dipeptide):
    seq_count = 0
    seqLen=len(specific_dipeptide)
    for i in range(len(sequence) - seqLen +1):
        seq = sequence[i:i+seqLen]
        if seq == specific_dipeptide:
            seq_count += 1
    return seq_count

def count_specific_xFxFG(sequence):
    seq_count = 0
    seqLen=5
    for i in range(len(sequence) - seqLen +1):
        seq = sequence[i:i+seqLen]
        if seq[1] == "F" and seq[3] == "F" and seq[4] == "G":
            seq_count += 1
    return seq_count

def sliding_window(sequence, window_size, specific_dipeptide):
    counts = []
    for i in range(len(sequence) - window_size + 1):
        window = sequence[i:i+window_size]
        dipeptide_count = 0
        for i in range(len(window) - 1):
            dipeptide = window[i:i+2]
            if dipeptide == specific_dipeptide:
                dipeptide_count += 1
        counts.append(dipeptide_count)
    return counts

file_path = 'Araport11_genes.201606.pep.fasta.gz'
names, sequences, genes = read_fasta(file_path)
window_size = 200

n=0
resultsFile = open('FG-Analysis-Araport11.txt', 'w')

print(f"#\tGene\tGene Name\tmax #FG in 200 window\t#FG\tLength\t#GLFG\t#xFxFG\tStart of Max Window", file = resultsFile)
for name, sequence, gene in zip(names, sequences, genes):
    resultFG = count_specific_sequence(sequence, "FG")
    if resultFG > 0 and len(sequence)>=200:
        resultGLFG = count_specific_sequence(sequence, "GLFG")
        resultxFxFG = count_specific_xFxFG(sequence)
        window_counts = sliding_window(sequence, window_size, "FG")
        index_of_max_value = window_counts.index(max(window_counts))
        print(f"{n}\t{name}\t{gene}\t{max(window_counts)}\t{resultFG}\t{len(sequence)}\t{resultGLFG}\t{resultxFxFG}\t{index_of_max_value}", file = resultsFile)
    n +=1
resultsFile.close()
```
